# Dissecting Complex Interactions Between Ferroptosis and the Proteasome

**DOI:** 10.64898/2026.03.28.714855

**Authors:** Magdalena B. Murray, Rishi Upadhyay, Krystina Szylo, Aastha Gautam, Judith Goncalves, Giovanni Forcina, Ananya Vinayak, Onn Brandmann, Scott J. Dixon

## Abstract

The proteasome is an essential multiprotein complex whose inhibition can lead to apoptosis. Ferroptosis is a non-apoptotic cell death mechanism whose fundamental regulation continues to be elucidated. How proteasome function regulates ferroptosis sensitivity is poorly understood and difficult to study given the essential nature of the proteasome. Here, we isolated the effects of proteasome inhibition on ferroptosis by combining direct cell death imaging, cell death pathway-specific inhibitors, and mathematical modeling. We find that proteasome inhibition enhances sensitivity to ferroptosis induced by glutathione peroxidase 4 (GPX4) inhibition while simultaneously promoting resistance to ferroptosis induced by system x_c_ ^-^ inhibition. Sensitization to GPX4 inhibition requires protein synthesis but not the apoptosis execution machinery and is opposed by the activating transcription factor 4 (ATF4) stress response pathway. This work demonstrates a complex role for proteasome function in ferroptosis regulation and establishes new methods to dissect cross-talk between ferroptosis and essential cellular processes.

## INTRODUCTION

Drug combinations are common in biology and a mainstay for cancer therapy. An underexplored facet of the drug combination problem concerns the interaction between agents that engage distinct cell death mechanisms. While most classic chemotherapy agents induce apoptosis^1^, new agents that trigger non-apoptotic modes of cell death, including ferroptosis, cuproptosis or disulfidptosis, are emerging for cancer therapy.^2–6^ Agents that induce different cell death modes may interact in unexpected ways, for example if one pathway is quicker to execute than another, or if the execution of one mechanism impinges directly on another.^7,8^ When perturbing essential processes it is important to disentangle the proximate effects of target protein inhibition from the distal induction of cell death that may result as a consequence. Understanding these complex relationships will help maximize the utility of drug treatments for cancer and other diseases.

Ferroptosis is a non-apoptotic form of cell death that is characterized by the iron-dependent accumulation of membrane lipid peroxides.^9,10^ Ferroptosis is normally prevented by the activity of several different inhibitory mechanisms.^11,12^ The selenoprotein glutathione peroxidase 4 (GPX4) restrains lethal lipid peroxidation and is likely the most important ferroptosis inhibitor in most cells.^13,14^ The small molecule 1*S*,3*R*-RSL3 (hereafter, RSL3) is a prototypic ferroptosis inducer that inhibits GPX4 and potentially other selenoproteins.^14–16^ The system x_c_^-^ antiporter erastin, which prevents the uptake of cystine, is another canonical ferroptosis inducer.^17^ Cystine, once reduced to cysteine inside the cell, is used to synthesize the GPX4 cofactor glutathione, along with other metabolites that help prevent ferroptosis.^18,19^ Ferroptosis is important for normal and disease biology^12^ and we are interested in understanding the fundamental regulation of this process.

The proteasome is a large molecular machine that catabolizes ubiquitinated proteins and is a target for cancer therapy, especially in multiple myeloma. Proteasome function may intersect with the ferroptosis network, but the literature in this area is complex.^8,17,20^ In hepatoma cells, the proteasome inhibitor bortezomib can induce lipid peroxidation^21^, a canonical feature of ferroptosis. Proteasome inhibition may also sensitize cells to ferroptosis inducers, possibly by affecting intracellular iron or glutathione levels.^21–24^ In triple negative breast cancer, inhibition of the proteasome and the epigenetic regulator bromodomain containing 4 (BRD4) is sufficient to induce ferroptosis.^25,26^ However, in other contexts, proteasome inhibition reduces ferroptosis sensitivity.^8^ The fact that proteasome inhibition itself causes apoptosis^27,28^ has the immediate potential to confound the analysis of ferroptosis if the appropriate controls and cell death detection methods are not employed.

Here, building on results from new CRISPR/Cas9 genetics screens, we examined how ferroptosis sensitivity was regulated by proteasome function. We induced ferroptosis and perturbed proteasome function mainly using combinations of small molecule inhibitors. We developed methods to isolate the proximate effects of proteasome-mediated protein turnover on ferroptosis, separate from the ability of proteasome inhibitors to trigger apoptosis. We find that proteasome inhibition enhances sensitivity to GPX4 inhibition while simultaneously reducing sensitivity to system x_c_ ^-^ inhibition. These results reinforce the context-dependent nature of ferroptosis regulation^29,30^ and highlight a complex role for proteasome function in ferroptosis regulation.

## RESULTS

### CRISPR screens identify stimulus-specific ferroptosis sensitizers

To identify ferroptosis-sensitizing genes, we performed parallel pooled CRISPR screens in populations of NALM6 leukemia cells harboring a doxycycline-inducible Cas9 transgene and transduced with a genome-wide sgRNA library^31^ (**Figure 1A**). NALM6 Cas9-expressing cells growing in suspension were treated with the vehicle control dimethyl sulfoxide (DMSO), RSL3 at 120-300 nM, or erastin2 at 150 nM. After eight days, cells treated with DMSO underwent 7.5 population doublings versus 5.9 doublings for both RSL3 and erastin2. Cells were harvested and analyzed by deep sequencing, revealing significantly depleted (sensitizer) or enriched (suppressor) genes (FDR q < 0.05). Different sensitizer and suppressor genes were recovered for RSL3 versus erastin2 (**Figure 1B-D**). For RSL3, sensitizer genes were involved in selenium uptake (*LRP8*)^32,33^, tetrahydrobiopterin synthesis (*GCH1*, *PTS*, *SPR*)^30,34^, lipoprotein uptake (*PAPSS1*)^35^, and vitamin K metabolism (*VKORC1L1*)^36^, while for erastin2 sensitizers included both components of system x_c_ ^-^ (*SLC7A11*, *SLC3A2*)^17^, and a different gene involved in selenium metabolism (*PRDX6*)^37–39^ (**Figure 1B,C**). While the screening conditions were biased toward identifying sensitizer genes, the lipid metabolic gene acyl-CoA synthetase long-chain family member 4 (*ACSL4*) was identified as a suppressor gene when using RSL3 but not erastin2, consistent with previous results^40^ (**Figure 1B,C**). The recovery of known ferroptosis regulators provided confidence in the quality and generalizability of these screening results.

**Figure 1.**
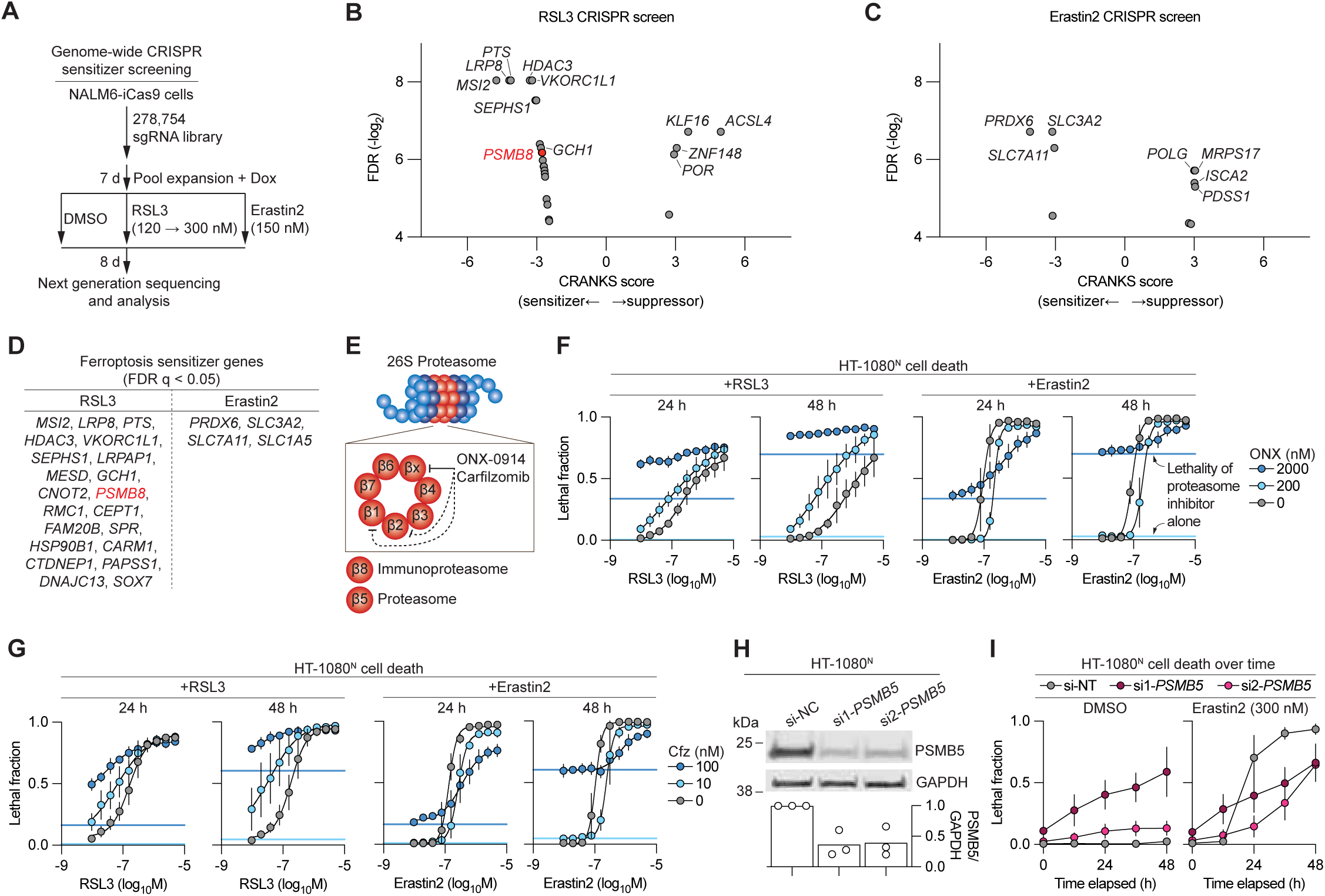
Proteasome inhibition modulates ferroptosis sensitivity. (A) Outline of CRISPR screens to identify ferroptosis regulators. (B) Results of the RSL3 CRISPR screen, with PSMB8 highlighted in red. (C) Results of the erastin2 CRISPR screen. (D) Summary of all significant (FDR q < 0.05) sensitizer genes identified in the CRISPR screens using RSL3 and erastin2, with *PSMB8* highlighted in red. (E) Cartoon of the normal proteasome and immunoproteasome with relevant subunits and inhibitors. (F) Cell death determined by imaging of live (nuclear mKate2-positive) and dead (SYTOX Green-positive) cells. Live and dead cell counts were integrated into the lethal fraction score (0 = all cells in the population alive, 1 = all cells in the population dead). The solid horizontal lines indicate the basal lethality of ONX-0914 (ONX) alone at different concentrations, color-coded as per the legend. Data are mean ± SD from two to three independent experiments. (G) Cell death determined by imaging. The solid horizontal lines indicate the basal lethality of carfilzomib (Cfz) alone at different concentrations, color-coded as per the legend. (H) Protein expression determined by immunoblotting. Blot image is representative of three independent experiments. Quantification of PSMB5 expression normalized to GAPDH expression from three independent experiments is shown below the blot image. (I) Cell death determined by imaging. si-NT, non-targeting (control) short interfering RNA. Data in **G** and **I** are mean ± SD from three independent experiments.

*PSMB8* was one of several underexplored RSL3-specific sensitizer genes identified in the CRISPR screen (**Figure 1B,D**). *PSMB8* encodes a subunit of the immunoproteasome, a variant proteasome linked to oxidative stress resistance (**Figure 1E**).^41,42^ *PSMB8* is broadly expressed in cancer cells, including many adherent cell lines commonly used for ferroptosis mechanistic studies (**Figure S1A**). We therefore pursued follow up studies in adherent cancer cell lines. HT-1080^N^ fibrosarcoma cells express the live-cell marker nuclear-localized mKate2 (denoted by the superscript ‘N’) and can be incubated with the dead cell dye SYTOX Green, enabling live and dead cells to be directly counted over time using imaging.^43^ To follow up on the results of the CRISPR screen, we initially treated HT-1080^N^ cells concurrently with a putative immunoproteasome inhibitor, ONX-0914 (ref.^44^) and RSL3 or erastin2, administered in 10-point, 2-fold dose response series. DMSO served as the vehicle control. Cell death at 24 h and 48 h was quantified by integrated counts of live and dead cells into the lethal fraction score, which represents population cell death on a 0 (all alive) to 1 (all dead) scale.^43^ Cells treated with ONX-0914 were sensitized to RSL3 and simultaneously more resistant to erastin2 (**Figure 1F**). For example, treatment with 200 nM ONX-0914 increased RSL3 potency (EC_50_) at 48 h by ∼10-fold compared to cells treated with DMSO (**Figure S1B**).

The immunoproteasome and constitutive proteasome share most of the same subunits, with a few substitutions including PSMB8 (immunoproteasome) for PSMB5 (constitutive).^42^ HT-1080 and other adherent cancer cells express *PSMB8*, but often express *PSMB5* at much higher levels (**Figure S1A**). In the DepMap database (25Q3)^45^, *PSMB5* was a genetic dependency for most tested cancer cell lines (1119 of 1186, 94%) while *PSMB8* was not (1 of 1186, > 0.1%). ONX-0914 can partially inhibit PSMB5 (ref.^46^) and was lethal to HT-1080^N^ cells alone. Thus, we suspected that ONX-01914 modulated ferroptosis by inhibiting PSMB5, not PSMB8. Indeed, the more specific PSMB8 inhibitor M3258 (ref.^47^) did not modulate ferroptosis in HT-1080^N^ cells (**Figure S1C**). Moreover, the potent covalent PSMB5 inhibitor carfilzomib^48^ sensitized HT-1080^N^ cells to RSL3 and promoted resistance to erastin2 at lower doses than ONX-01914 (**Figure 1G**).

To test whether constitutive proteasome inhibition was sufficient to modulate ferroptosis sensitivity, we silenced *PSMB5* expression using two independent short interfering RNAs (siRNAs). In HT-1080^N^ cells, silencing of *PSMB5* reduced PSMB5 protein levels, was partially toxic alone, sensitized to a low dose of RSL3 (50 nM), and conferred resistance to a lethal dose of erastin2 (300 nM), consistent with results obtained using carfilzomib (**Figure 1I,H S1D**). RSL3 is a covalent inhibitor that likely has off-target effects, especially at high doses.^49^ However, carfilzomib sensitized HT-1080^N^ cells to two other GPX4 inhibitors, ML162 and Compound 28 (ref.^50^) (**Figure S1E,F**). Furthermore, when treated with carfilzomib, clonal HT-1080^N^ *GPX4*^KO^ cell lines^51^ were more sensitive to the withdrawal of ferrostatin-1, a ferroptosis-specific radical trapping antioxidant^9^, than a matched Control cell line, implying greater ferroptotic sensitivity (**Figure S1G**,**H**).

To examine whether erastin2 was acting in an on-target manner on system x_c_ ^-^, we directly deprived U-2 OS^N^ osteosarcoma cells of cystine in the cell culture medium. Carfilzomib treatment (10 nM) protected U-2 OS^N^ cells from cell death induced by cystine depletion, with this effect being clearest when background carfilzomib-induced apoptosis was suppressed by cotreatment with the pan-caspase inhibitor Q-VD-OPh (**Figure S1I**). Carfilzomib treatment did not generally alter sensitivity to cell death induced by a diverse panel of non-ferroptotic lethal agents, suggesting a degree of specificity to the observed effects with RSL3 and erastin2 (**Figure S1J**). Altogether, these data suggested that proteasome inhibition modulated ferroptosis in a stimulus-specific manner.

### Ferroptosis inducers do not directly inhibit the proteasome

How proteasome inhibitors altered ferroptosis sensitivity was unclear. We first examined whether ferroptosis inducers directly modulated proteasome function. Three lines of evidence suggested that this was unlikely. First, in HT-1080 and 293T cells expressing a destabilized ornithine decarboxylase-GFP reporter of proteasome activity^52^, RSL3 treatment caused only weak reporter stabilization compared to carfilzomib and bortezomib, while erastin2 had no effect (**Figure 2A, S2A**). RSL3-induced reporter stabilization was suppressed in cells cotreated with ferrostatin-1, indicating that this was likely caused by lipid peroxidation, not direct proteasome inhibition (**Figure 2A, S2A**). Second, in permeabilized HT-1080^N^ cells, RSL3 and erastin2 did not strongly inhibit proteasome-associated chymotrypsin (β5)-like, trypsin (β2)-like, or caspase (β1)-like activity measured using fluorogenic substrates (**Figure 2B**). Third, RSL3 and erastin did not stabilize NFE2 like bZIP transcription factor 1 (NFE2L1), a protein that is normally rapidly turned over by proteasome-mediated degradation (**Figure S2B,C**).^53,54^

**Figure 2.**
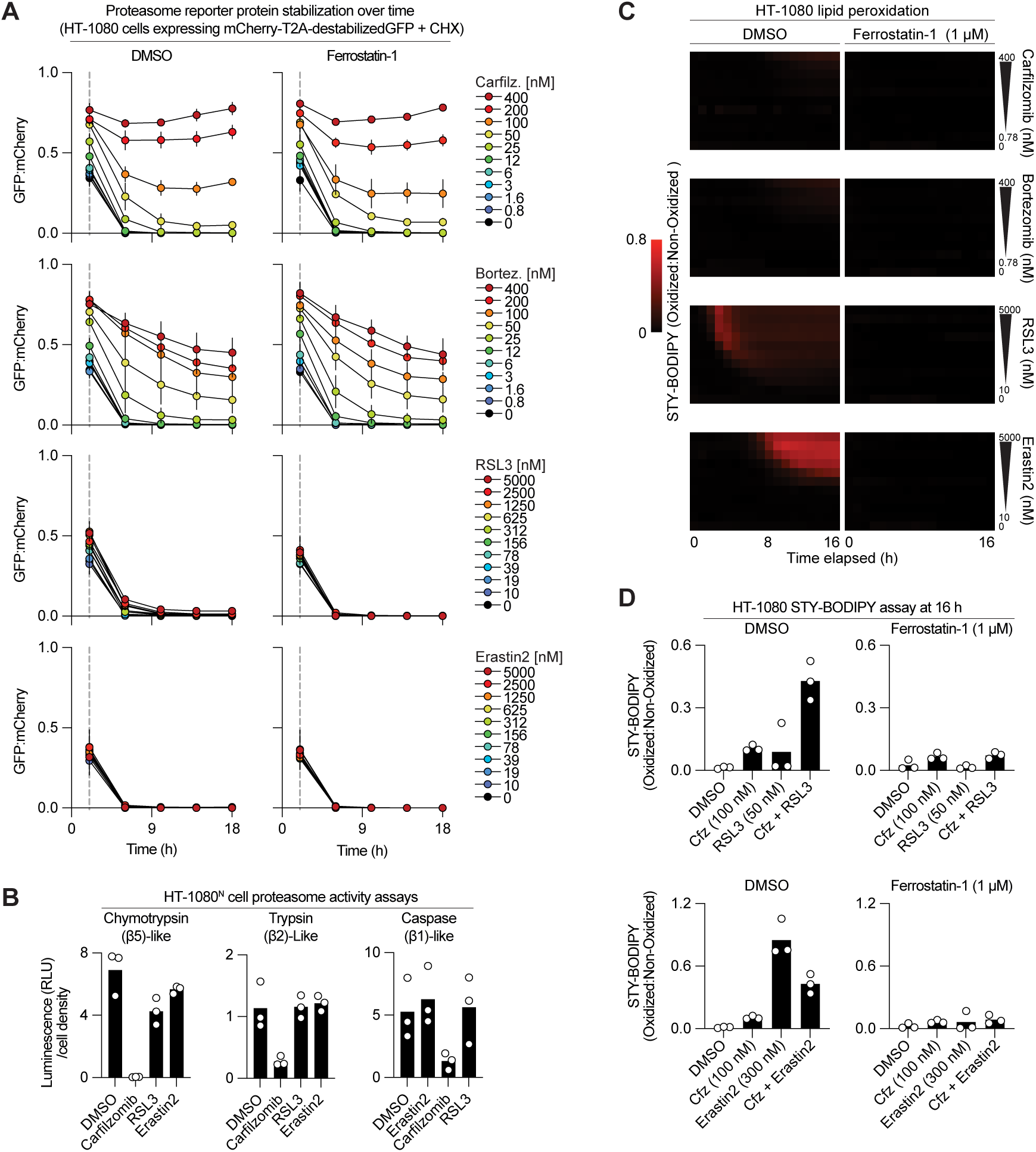
Proteasome inhibition is not a direct ferroptosis trigger. (A) Proteasome reporter assay. Cells were treated with inhibitors and cycloheximide for 2 h, then cycloheximide was washed out and cells were imaged to assess reporter protein stability. Carfilz, carfilzomib; Bortez, bortezomib. (B) Biochemical analysis of proteasome enzymatic activity. (C) Analysis of lipid peroxidation over time using STY-BODIPY. (D) Analysis of lipid peroxidation over time using STY-BODIPY. Cfz, carfilzomib. Data in **A** and **C** are mean ± SD from three or four independent experiments. Data in **B** and **D** are individual datapoints from three independent experiments

We next tested mechanisms reported in the literature which could directly link proteasome inhibition to ferroptosis regulation. NFE2L1 stabilization upon proteasome inhibition might promote ferroptosis resistance.^20,51^ However, carfilzomib pretreatment (100 nM, 4 h) sensitized to RSL3 and suppressed erastin2-induced cell death to a similar extent in HT-1080^N^ Control and gene-disrupted *NFE2L1*^KO^ cell lines generated using CRISPR/Cas9 technology (**Figure S2D**). Proteasome inhibition may directly increase lipid peroxidation^21,55,56^ which could synergize with ferroptosis inducers. We developed a high-throughput time-lapse lipid peroxidation monitoring assay based on the chemical probe STY-BODIPY^57^ and found that, unlike RSL3 and erastin2, carfilzomib and bortezomib did not induce substantial probe oxidation (**Figure 2C**). Under some conditions, proteasome inhibitors can deplete cells of the anti-ferroptotic amino acid cysteine.^58^ However, in our standard HT-1080^N^ cell culture conditions, proteasome inhibitors did not cause acute depletion of cysteine or other amino acids and proteasome inhibitor-induced cell death was not suppressed by supplementing cells with *N*-acetyl-cysteine^9^ (**Figure S2E,F**).

Given these results, it seemed likely that proteasome inhibition altered ferroptosis indirectly. Consistent with this possibility, carfilzomib enhanced STY-BODIPY oxidation in cells cotreated with RSL3 and suppressed STY-BODIPY oxidation in cells cotreated with erastin2, and these effects were rescued by ferrostatin-1 (**Figure 2D**, note that the signal observed with carfilzomib alone was ferrostatin-1-insensitive and likely non-specific). We also confirmed that cell death induced by carfilzomib and bortezomib as single agents was not suppressed by ferrostatin-1 (**Figure S2G**,**H**). We inferred that proteasome inhibitors modulated ferroptosis by altering the abundance of one or more mediators that are depleted or accumulated as a consequence of proteasome inhibition.

### High-throughput lethal compound interaction profiling

Thus far, our results suggested that proteasome inhibitors modulated sensitivity to ferroptosis-inducing agents indirectly. However, investigating these compound-compound interactions was difficult: proteasome inhibitors cause apoptosis^27,28^ (**Figure S2G,H**) and in cells dying by apoptosis it can be hard to see the effect proteasome inhibition *per se* on ferroptosis. It also seemed likely that the degree and timing of proteasome inhibition could alter the magnitude of ferroptosis modulation. To address these challenges, we developed time-staggered kinetic modulatory profiling (tskMP). TskMP incorporates graded functional perturbation, mechanism-specific cell death inhibitors, time-lapse cell death imaging, and mathematical modeling to assess interactions between two independently lethal agents.

For our tskMP analysis, we first generated different degrees of proteasome inhibition in HT-1080^N^ and U-2 OS^N^ cells by treating with carfilzomib at 1, 10 or 100 nM, each for 0, 1, 4, or 24 h (**Figure 3A**). These doses of carfilzomib produced a graded and relatively specific inhibition of proteasomal chymotrypsin (β5)-like activity (**Figure S3A**). To these pretreated cells we then added erastin2 or RSL3, each in a 10-point, two-fold dose response series. To isolate different lethal mechanisms, these treatments were performed in parallel plates using cells cotreated with vehicle (DMSO), the ferroptosis inhibitor ferrostatin-1 (1 µM), or the pan-caspase inhibitor Q-VD-OPh (25 µM). Following compound addition, cell death was monitored every 6 h for 48 h by time-lapse imaging. The experiment was repeated three times and the resulting dataset contained ∼28,000 population-level cell death measurements. Compound combination effects were evaluated using an established method based on an adaptation of the Bliss model of compound interactions to kinetic data.^8^ Heatmaps were used to summarize deviation (D) in the observed amount of cell death for each compound combination from the expected amount based on single agent activities, with positive D scores indicating synergy and negative D scores indicating antagonistic interactions (**Figure 3B**,**C**).

**Figure 3.**
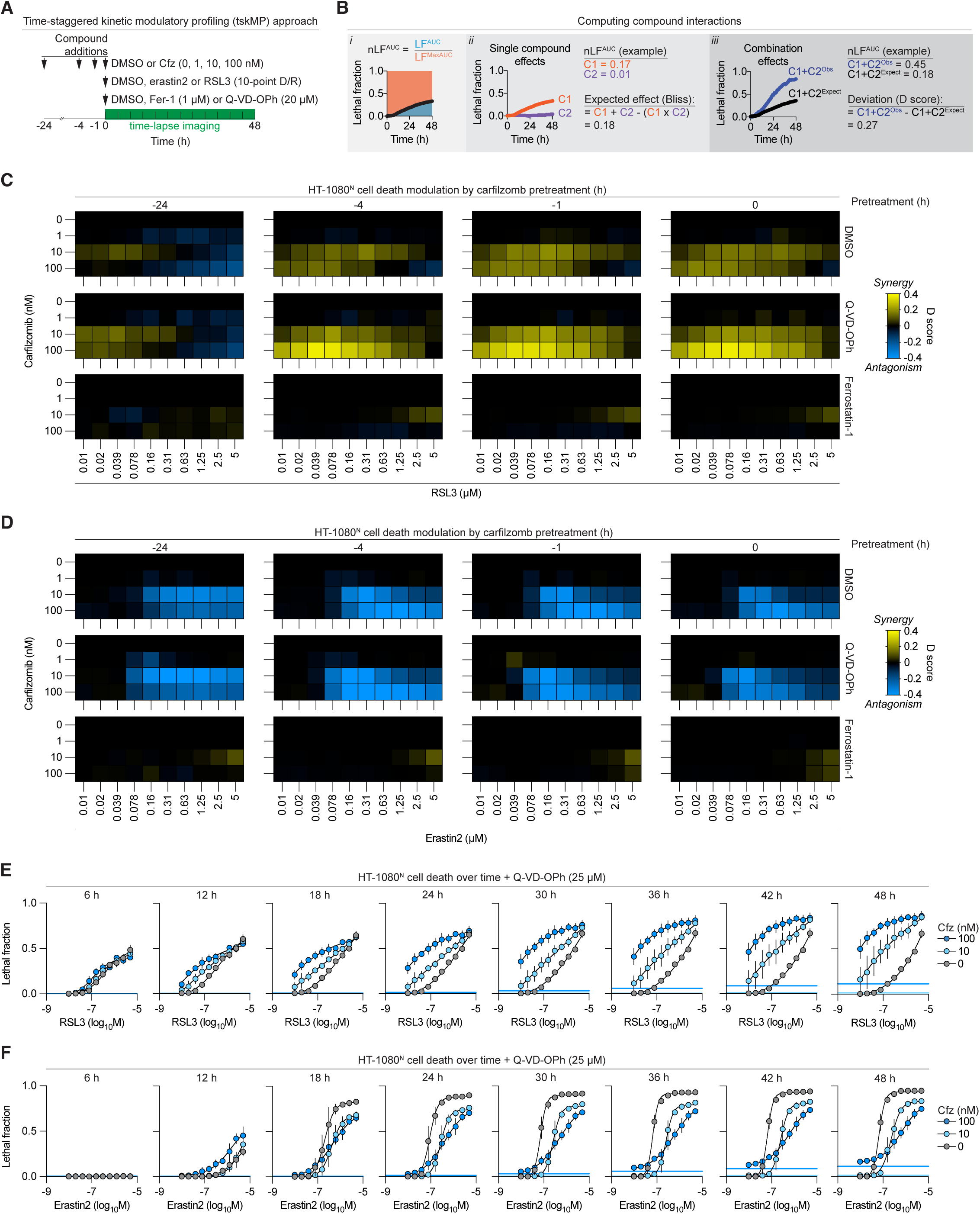
A time-staggered kinetic modulatory profile analysis. (A) Outline of the time-staggered kinetic modulatory profile (tskMP) screen experimental design. (B) Outline of the method to analyze compound-compound interactions. *i*, the area under the curve (AUC) of lethal fraction scores over time is normalized to the total potential lethal fraction AUC. *ii*, The expected effect of combining both compounds together is determined using the Bliss equation. Iii, The observed (Obs) effect of the compound combination is subtracted from the expected effect (Expect), yielding a deviation (D) score. Each step is illustrated by example data. (C, D) Summary of compound-compound deviation (D) scores. Ferrostatin-1 was used at 1 µM, Q-VD-OPh at 25 µM. Data are mean values from three independent experiments. (E,F) Deconvoluted kinetic data from the tskMP analysis in panels C and D. Cell death determined by imaging. The solid horizontal lines indicate the basal lethality of carfilzomib alone, color-coded as per the legend. Data are mean ± SD from three independent experiments.

In HT-1080^N^ cells pretreated with 10 nM or 100 nM carfilzomib for 0 to 4 h we observed clear synergy across a range of RSL3 doses (**Figure 3C**). Interestingly, longer carfilzomib pretreatment was not associated with greater RSL3 synergy, possibly suggesting that this synergy arises only once GPX4 inhibition has been initiated. Pretreatment with 10 nM carfilzomib for 24 h also yielded synergy with low doses of RSL3 (**Figure 3C**). We suspected that synergy between 100 nM carfilzomib and RSL3 under these same conditions was masked by carfilzomib-induced apoptosis. Indeed, when cells were cotreated with Q-VD-OPh, we readily detected synergy between 100 nM carfilzomib and low doses of RSL3 (**Figure 3C**). In the 4 and 24 h carfilzomib pretreatment conditions, we observed apparent antagonistic interactions between carfilzomib and higher doses of RSL3 (negative D scores) (**Figure 3C**). Once again, cotreatment with Q-VD-OPh revealed evidence of synergy between carfilzomib and RSL3 under these conditions (**Figure 3C**). This synergy was eliminated in cells co-treated with ferrostatin-1, indicating that this effect was due to increased ferroptosis not a distinct mode of cell death (**Figure 3C**). Similar patterns of synergistic interactions between carfilzomib and RSL3 were also observed in U-2 OS^N^ (**Figure S3B**).

For erastin2, a distinct pattern of compound interactions was observed in the tskMP analysis. Consistent with earlier data, pretreatment with carfilzomib (10 nM or 100 nM) for 24, 4, 1 or 0 h antagonized erastin2-induced cell death, but for 10 nM carfilzomib these effects were strongest with longer pretreatments (e.g., 24 h versus 0 h) (**Figure 3D**). Cotreatment with Q-VD-OPh helped reveal the ability of carfilzomib to antagonize erastin2-induced ferroptosis, especially at low doses of erastin2 (**Figure 3D**). Cell death in cells treated with carfilzomib and erastin2 was also blocked by ferrostatin-1 (**Figure 3D**). In U-2 OS^N^ cells, carfilzomib (10 and 100 nM) likewise antagonized erastin2 lethality, albeit with weaker effects across erastin2 doses and pretreatment durations (**Figure S3C**). This tskMP analysis suggested that proteasome inhibition consistently modulated ferroptosis sensitivity across a range of conditions, with the timing of proteasome inhibition relative to ferroptosis induction being more relevant to erastin2-induced ferroptosis suppression than RSL3-induced ferroptosis sensitization.

In the tskMP analyses, Q-VD-OPh cotreatment made it easier to detect synergy between carfilzomib and RSL3 and antagonism between carfilzomib and erastin2 (**Figure 3C,D**). We examined how compound synergy and antagonism manifested over time in Q-VD-OPh-cotreated HT-1080^N^ cells. For simplicity, we focused on the 0 h carfilzomib pretreatment condition. RSL3 was lethal within 6 h but the addition of carfilzomib did not enhance cell death until 12 h, with synergy increasing thereafter until the end of the observation period (**Figure 3E**). For erastin2, the addition of carfilzomib also initially appeared to increase cell death at 12 h, but thereafter between 18 and 48 h, cell death increased more rapidly in cells treated with erastin2 alone than in cells cotreated with either 10 nM or 100 nM carfilzomib (**Figure 3F**). Thus, using tskMP analysis and cell death-mechanism-specific inhibitors, we could resolve complex interactions proteasome inhibition and ferroptosis induction.

### Apoptosis execution does not modulate ferroptosis

Proteasome inhibition appeared to modulate ferroptosis sensitivity in a complex manner depending on the ferroptosis-inducing stimulus. These phenotypes become stronger or easier to detect in cells co-treated with the pan-caspase inhibitor Q-VD-OPh. In some contexts, caspase inhibition can shift the mode of cell death in response to a lethal stimulus from one mechanism to another.^59^ We therefore examined whether cells treated with carfilzimib and Q-VD-OPh responded to RSL3 and erastin2 by undergoing ferroptosis or another form of cell death. We observed that in cells treated with carfilzomib and Q-VD-OPh, the cell death caused by RSL3 and erastin2 was fully suppressed by ferrostatin-1, indicating that caspase inhibition was not switching the mode of cell death in our combination treatments (**Figure S4A**).

Proteasome inhibition ultimately results in the induction of apoptosis.^25,26^ An important question at this stage was whether proteasome inhibitors modulated ferroptosis by engaging the apoptosis cascade. Carfilzomib modulated the lethality of RSL3 and erastin2 in HT-1080^N^ and U-2 OS^N^ when Q-VD-OPh was present, indicating that caspase activity was not important for these effects (**Figure 3C-H**). However, upstream events in the apoptosis cascade, including MOMP and cytochrome c release, are reported to sensitize^60^ or lead to resistance^61^ to ferroptosis. To investigate whether these events were needed for proteasome inhibition to modulate ferroptosis, we first overexpressed MCL1, a BCL2-family protein that inhibits BAX/BAK1-dependent MOMP (**Figure 4A,B**).^62^ MCL1-overexpressing cells resisted carfilzomib treatment, as expected, yet carfilzomib synergized with RSL3 and antagonized erastin2 to a similar extent as in cells expressing the empty vector (EV) control, similar to the effects of Q-VD-O-Ph (**Figure 4C**).

**Figure 4.**
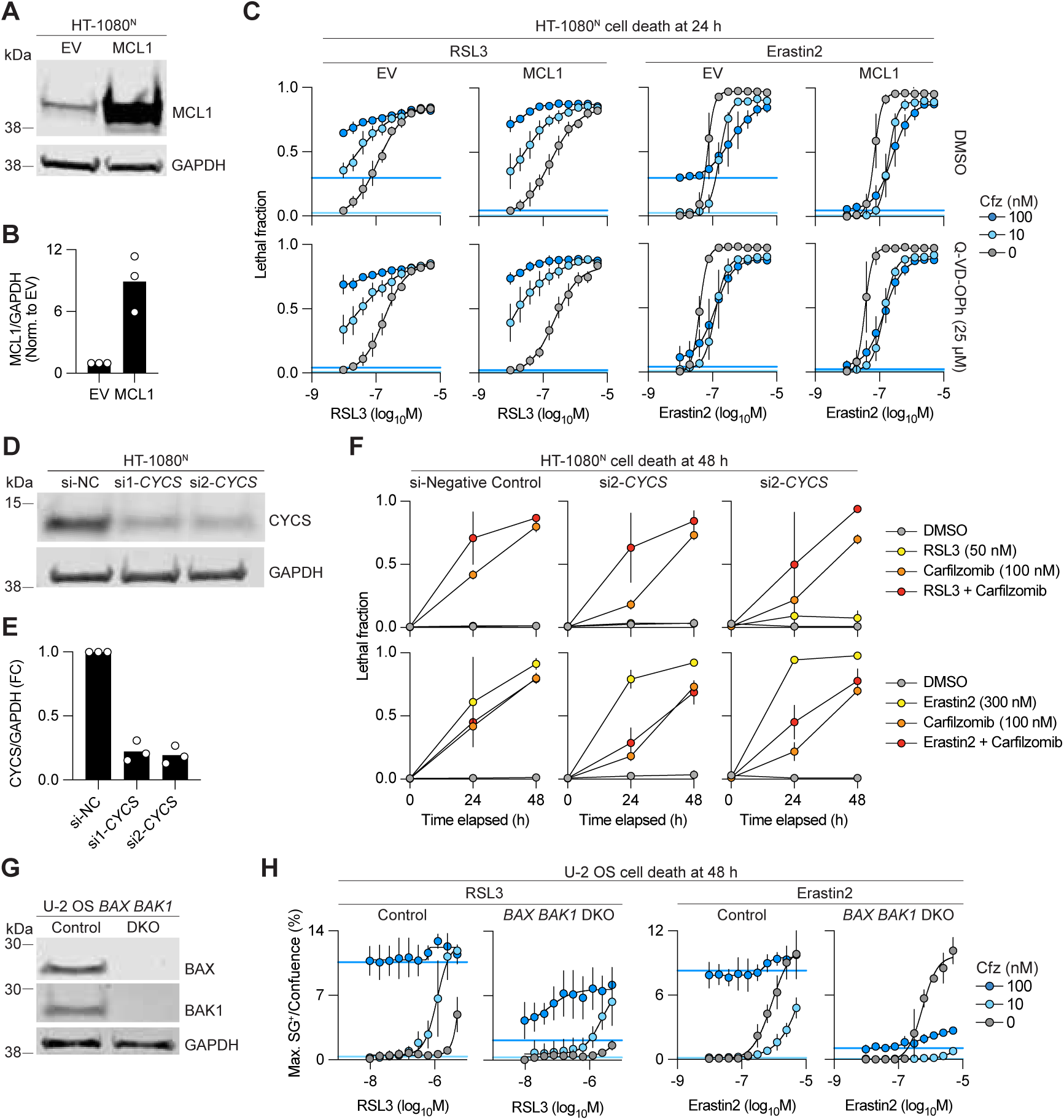
Ferroptosis modulation is independent of apoptosis. (A) Protein abundance determined by immunoblotting. (B) Quantification of relative MCL1 protein abundance from **A**. (C) Cell death determined by imaging of live (nuclear mKate2-positive) and dead (SYTOX Green-positive) cells. Live and dead cell counts were integrated into the lethal fraction score (0 = all cells in the population alive, 1 = all cells in the population dead). The solid horizontal lines indicate the basal lethality of carfilzomib (Cfz) alone at different concentrations, color-coded as per the legend. (D) Protein abundance determined by immunoblotting. (E) Quantification of relative CYCS protein abundance from **D**. (F) Cell death determined by imaging. (G) Protein abundance determined by immunoblotting. (H) Cell death determined by imaging. Blots in **A**, **D** and **G** are representative of three independent experiments. Data in **C**, **F** and **H** are mean ± SD from three to four independent experiments

Next, we used siRNA to directly silence expression of *CYCS* (encoding cytochrome c) (**Figure 4D,E**). Partial silencing of *CYCS* inhibited basal carfilzomib-induced (i.e., apoptotic) cell death at 24 h but did not prevent carfilzomib from sensitizing to RSL3 or promoting resistance to erastin2 (**Figure 4F**). We also performed experiments in U-2 OS control and *BAX BAK1* double knockout (DKO) cells^63^ and found that carfilzomib alone was less lethal in DKO cells, as expected for a pro-apoptotic agent, yet still synergized with RSL3 and antagonized the lethality of erastin2 (**Figure 4G,H**). Mathematically, strong synergy (D) scores were observed between carfilzomib and RSL3 or erastin2 in both control and DKO cells, with these effects being easier to discern in DKO cells (**Figure S4B**).

We also tested whether engaging the apoptosis machinery using a distinct trigger would modulate ferroptosis in HT-1080^N^ cells in the same manner as proteasome inhibition. For these experiments we selected the topoisomerase inhibitor camptothecin. Like carfilzomib, camptothecin induced substantial basal cell death in HT-1080^N^ cells that was blocked by cotreatment with Q-VD-OPh, consistent with the induction of apoptosis (**Figure S4C**). In conditions where apoptosis execution was blocked using Q-VD-OPh, we observed that 10 nM carfilzomib increased the potency (EC_50_) of RSL3 by over 40-fold, whereas a roughly equivalently lethal dose of camptothecin increased the potency of RSL3 by only 2-fold (**Figure S4D**). Thus, proteasome inhibition seemed more likely to modulate ferroptosis through proximate effects on protein turnover rather than by engaging the core apoptotic cascade.

### Protein synthesis is necessary for sensitization to GPX4 inhibition

Proteasome inhibition can stabilize wild-type p53.^25^ We previously showed that stabilizing wild-type p53 by inhibiting mouse double minute 2 (MDM2) sensitized to GPX4 inhibitors and conferred resistance to system x_c_ ^-^ inhibitors^50,64^, the same pattern observed here with proteasome inhibitors. HT-1080^N^ and U-2 OS^N^ express wild-type p53. We therefore examined the effects of proteasome inhibition in cells lacking functional p53, finding that carfilzomib potently synergized with RSL3 in p53 null H1299^N^ non-small cell lung carcinoma cells and p53 mutant T98G^N^ glioblastoma cells, effects that were strongest in cells cotreated with Q-VD-OPh (**Figure S5A**,**B**). Carfilzomib also antagonized erastin2-induced cell death in T98G^N^ cells but not H1299^N^ cells (**Figure S5A,B**).

In addition to stabilizing proteins like p53, proteasome inhibition can also stimulate the expression of stress response proteins.^65–67^ We reasoned that by inhibiting protein synthesis, we could isolate the role of protein synthesis triggered by proteasome inhibition. Towards this end, we treated HT-1080^N^ cells with carfilzomib and the translation elongation inhibitor cycloheximide. Cycloheximide minimally affected the lethality of carfilzomib alone, suggesting that protein synthesis was not essential for proteasome inhibitors to cause apoptosis (**Figure 5A,B**). Cycloheximide alone suppressed cell death in response to erastin2, as observed previously, possibly because this spares existing intracellular cysteine (**Figure 5A**).^8,68^ Remarkably, cycloheximide completely abolished the synergy observed between carfilzomib and RSL3 (**Figure 5B,C**). Cycloheximide had similar effects on RSL3-induced cell death in H1299^N^ cells, consistent with a generalizable effect that was independent of p53 (**Figure S5C**). A distinct protein synthesis inhibitor, emetine, likewise yielded similar results in HT-1080^N^ cells (**Figure S5D**). Thus, protein synthesis inhibitors eliminated the synergy between proteasome inhibition and GPX4 inhibition.

**Figure 5.**
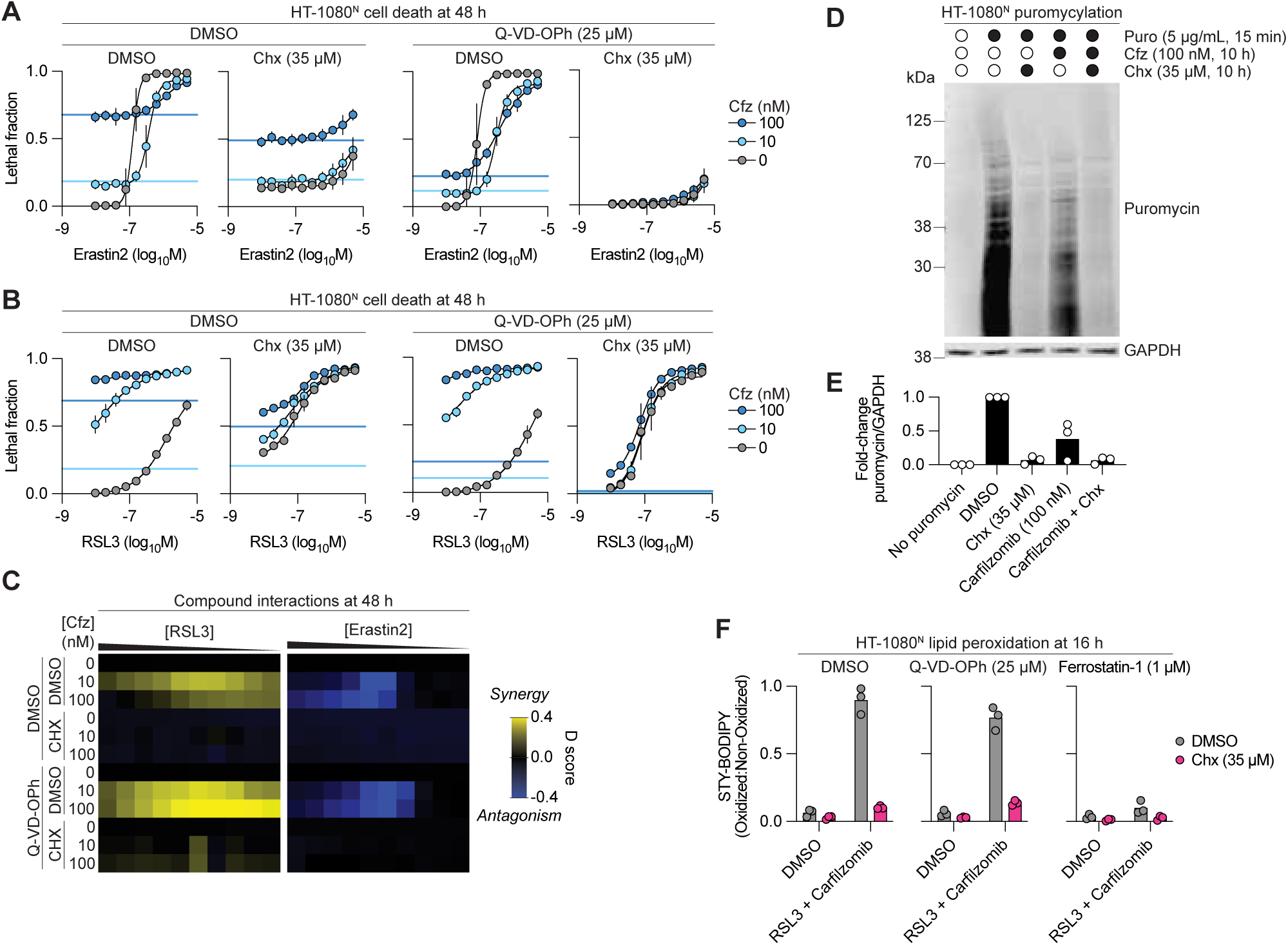
Ferroptosis modulation is dependent on protein synthesis. (A) Cell death determined by imaging of live (nuclear mKate2-positive) and dead (SYTOX Green-positive) cells. Live and dead cell counts were integrated into the lethal fraction score (0 = all cells in the population alive, 1 = all cells in the population dead). The solid horizontal lines indicate the basal lethality of carfilzomib (Cfz) alone at different concentrations, color-coded as per the legend. Cfz, carfilzomib; Chx, cycloheximide. (B) Cell death determined by imaging. (C) Summary of compound-compound deviation (D) scores. Heatmaps represent mean values from three independent experiments. (D) Total puromycin protein labeling determined by immunoblotting. Representative of three independent blots. (E) Quantification of C depicting relative puromycin abundance normalized to the DMSO condition. (F) Analysis of lipid peroxidation using STY-BODIPY. Data in **A** and **B** are mean ± SD from three to four independent experiments. Data in **E** and **F** show individual datapoints from three independent experiments.

Mechanistically, we confirmed using anti-puromycylation western blots^69^ that cycloheximide blocked protein synthesis on its own and in cells cotreated with carfilzomib (**Figure 5D**,**E**). We used our high-throughput STY-BODIPY imaging approach to examine the impact of protein synthesis inhibition on lipid peroxidation. The combination of carfilzomib (100 nM) and RSL3 (50 nM) resulted in substantial STY-BODIPY probe oxidation that was potently suppressed by cycloheximide or ferrostatin-1 (**Figure 5F**). Proteasome inhibition has been found to enhance^70,71^ or deplete^56^ intracellular glutathione. In HT-1080^N^ cells, carfilzomib treatment decreased intracellular total glutathione abundance, and this was prevented by cotreatment with cycloheximide (**Figure S5E**). Thus, protein synthesis appeared essential for synergy between proteasome blockade and GPX4 inhibition.

### ATF4 expression opposes the effects of proteasome inhibitors

Our results suggested that proteasome inhibition might sensitize to GPX4 inhibition by upregulating one or more proteins. To investigate, we performed nanoLC-MS-based proteome profiling of HT-1080^N^ and U-2 OS^N^ cells treated with vehicle (DMSO) or carfilzomib (100 nM), with cells harvested right before the onset of cell death at 15 or 22 h, respectively. Our results with cycloheximide and emetine suggested that protein synthesis following proteasome inhibition was critical for GPX4 inhibitor sensitization. Accordingly, we focused on 50 proteins that were undetectable in both cell lines basally and increased to detectable levels in response to proteasome inhibition, suggesting stress-induced synthesis (**Figure 6A**). Analyzing these results using the STRING database we discovered a highly interconnected set of proteins centered on activating transcription factor 4 (ATF4), whose increased abundance is a known response to proteasome inhibition^72^, including the ATF4 target glutathione-specific gamma-glutamylcyclotransferase 1 (CHAC1), the unfolded protein response (UPR) regulator DNA damage inducible transcript 3 (DDIT3, CHOP), and other proteins involved in the UPR (**Figure 6B**-**D**). ATF4 and its targets appear to either sensitize^73–75^ or suppress^76–78^ ferroptosis. Induction of ATF4 in response to proteasome inhibitor treatment was suppressed in cells cotreated with cycloheximide, consistent with expectations for a death-regulating protein (**Figure S6A**,**B**). Of note, cotreatment with carfilzomib and cycloheximide did not alter GPX4 levels or the ability of RSL3 to engage GPX4 covalently, as assessed by slower migration^50^ in polyacrylamide gels (**Figure S6A**,**B**).

**Figure 6.**
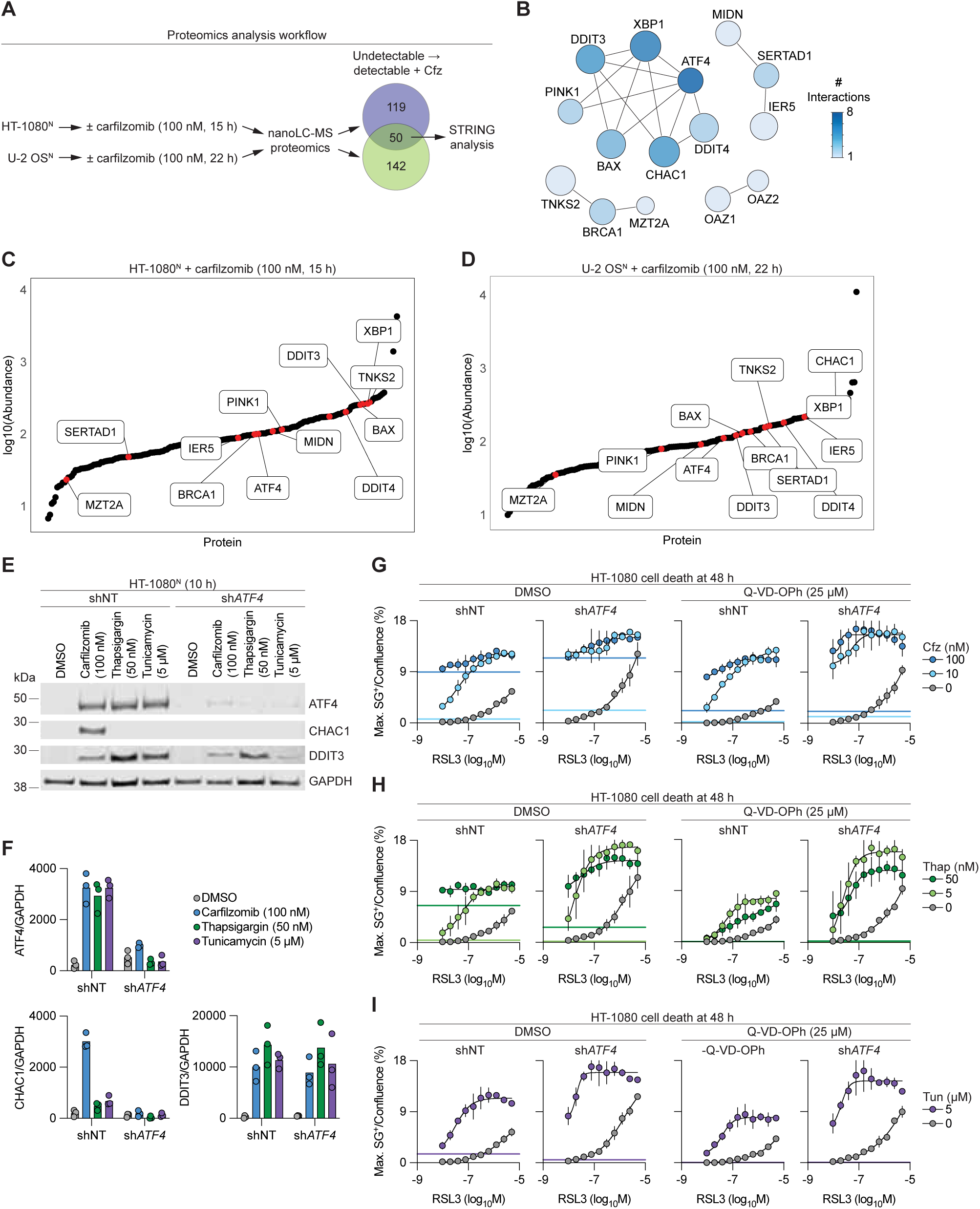
ATF4 negatively regulates ferroptosis. (A) Overview of proteomics and analysis workflows. Proteomics data are mean from four independent experiments. (B) Analysis of common proteins upregulated by carfilzomib treatment using the STRING database. (C) Absolute average increase in abundance following carfilzomib treatment. Proteins identified in STRING analysis indicated in red. (D) Absolute average increase in abundance following carfilzomib treatment. Proteins identified in STRING analysis indicated in red. (E) Protein abundance determined by immunoblotting. Blots are representative of three independent experiments. (F) Quantification of relative protein abundance from E. (G) Cell death determined by imaging of live (nuclear mKate2-positive) and dead (SYTOX Green-positive) cells. Live and dead cell counts were integrated into the lethal fraction score (0 = all cells in the population alive, 1 = all cells in the population dead). The solid horizontal lines indicate the basal lethality of carfilzomib (Cfz) alone at different concentrations, color-coded as per the legend. Cfz, carfilzomib. (H) Cell death determined by imaging. (I) Cell death determined by imaging. Data in **G**, **H** and **I** are mean ± SD from three independent experiments.

We reasoned that ATF4 could potentially decrease intracellular total glutathione and sensitize to GPX4 inhibition by upregulating CHAC1, a glutathione-degrading enzyme.^79,80^ If so, silencing of *ATF4* or *CHAC1* should prevent carfilzomib from sensitizing to RSL3. To test this hypothesis we first used HT-1080 cells stably expressing a short hairpin RNA (shRNA) against *ATF4* or a non-targeting (NT) control.^81^ Carfilzomib treatment resulted in the accumulation of ATF4, CHAC1 and DDIT3 in shNT cells and this accumulation of ATF4 and CHAC1, but not DDIT3, was blunted in sh*ATF4* cells (**Figure 6E**,**F**). However, with respect to cell death, sh*ATF4* cells exposed to carfilzomib exhibited greater synergy with RSL3 than observed in shNT cells, an effect best observed using Q-VD-OPh cotreatment (**Figure 6G, S6C**). Furthermore, direct CRISPR-mediated disruption of *CHAC1* in H1299 cells using established reagents^82^ did not prevent carfilzomib from sensitizing to RSL3 (**Figure S6D-F**). Thus, proteasome inhibitor-induced ATF4 expression appeared to promote GPX4 inhibitor resistance.

### Distinct inducers of the UPR sensitize to GPX4 inhibition

Proteasome inhibition is one of several stresses that can induce the unfolded protein response (UPR). To investigate whether the phenotypes observed using proteasome inhibition were generalizable to mechanistically distinct UPR inducers, we used the sarcoendoplasmic reticulum calcium ATPase (SERCA) inhibitor thapsigargin and the protein O-glycosylation inhibitor tunicamycin. In shNT cells, thapsigargin and tunicamycin both induced strong accumulation of ATF4 and DDIT3, but not CHAC1, distinguishing these stressors from proteasome inhibition (**Figure 6E**,**F**). Like with carfilzomib, cotreatment with thapsigargin or tunicamycin sensitized shNT cells to RSL3-induced cell death (**Figure 6H, S6C**). Likewise, genetic silencing of *ATF4* enhanced the synergy of RSL3 with thapsigargin or tunicamycin (**Figure 6H, S6C**), and these synergistic interactions were antagonized by cotreatment with emetine (**Figure S6G**-**I**). Thus, diverse UPR inducers sensitize to GPX4 inhibitor-induced ferroptosis and this is opposed by protein synthesis inhibition and the ATF4-mediated stress response.

## DISCUSSION

Our studies illustrate the challenge associated with understanding how an essential process, like proteasome-mediated protein turnover, regulates ferroptosis. Inhibiting proteasome function triggers apoptosis as a downstream consequence. To dissociate the upstream consequences of proteasome inhibition in ferroptosis regulation from the downstream induction of apoptosis we used a combination of high-throughput time-lapse cell death imaging, pathway-specific inhibitors, and mathematical analysis of compound interactions. Dissecting ferroptosis regulatory mechanisms that involve other essential processes could be undertaken using the same combination of approaches introduced here.

We initially identified the immunoproteasome subunit *PSMB8* as an RSL3-specific sensitizer gene in our NALM6 genome-wide CRISPR screen. However, in other cell lines, a PSMB8-specific inhibitor had no effect on ferroptosis, whereas PSMB5-binding inhibitors consistently modulated ferroptosis sensitivity across multiple models. Thus, ferroptosis it appears that ferroptosis can be modulated by inhibition of either the immunoproteasome or the constitutive proteasome. We speculate that, compared to HT-1080, U-2 OS and other adherent cells growing in standard culture, the NALM6 cells used in the CRISPR screen are more dependent upon the immunoproteasome than the constitutive proteasome for homeostatic protein turnover.

The execution of apoptosis, especially the induction of MOMP and release of cytochrome c, is reported to modulate ferroptosis either positively or negatively.^60,61^ However, using the techniques developed here to isolate apoptosis execution, we find that proteasome inhibition regulates ferroptosis independent of the apoptosis execution, including BAX/BAK1, MOMP, cytochrome c, and caspase activity. Our methods may have allowed us to ascertain a disconnect between apoptosis and ferroptosis that is hard to detect using traditional methods. Alternately, it is possible that cross-talk between the apoptosis execution machinery and ferroptosis is cell-type specific or only occurs in response to certain apoptosis-inducing conditions.

How does proteasome inhibition modulate ferroptosis distinctly depending on whether this process is induced by direct GPX4 inhibition or system x_c_ ^-^ inhibition? The proteasome regulates the turnover of many proteins; proteasome inhibition will allow for these proteins to accumulate. Two candidates that accumulate upon proteasome inhibition and have been connected to ferroptosis regulation, NFE2L1 (ref.^20,51^) and p53 (ref.^64^), were not required for the phenotypes observed here. The antioxidant transcription factor NFE2 like bZIP transcription factor 2 (NFE2L2, NRF2) is another proteasome-regulated ferroptosis suppressor.^83^ Accumulation of NRF2 upon proteasome inhibition might explain reduced sensitivity to system x_c_ ^-^ inhibition, but seems unlikely to explain enhanced sensitivity to GPX4 inhibitors.

Indeed, it is remarkable that proteasome inhibition synergizes with GPX4 inhibitors despite the accumulation of ATF4, NRF2 and potentially other ferroptosis negative regulators. How this occurs is not entirely clear. GPX4 is ubiquitinated and subject to proteasome-mediated turnover.^84^ It is possible that proteasome inhibition blocks new GPX4 protein synthesis and results in the accumulation of ubiquitinated GPX4 that is partly inactive or more susceptible to GPX4 inhibitor binding. This model would not, however, account for the importance of protein synthesis in the GPX4 sensitizing phenotypes observed here. Instead, we propose that the increased synthesis or one or more specific proteins upon proteasome inhibition drives cells into a GPX4 inhibitor-sensitive state.

Protein synthesis in the endoplasmic reticulum requires glutathione for protein folding^85^, and lower levels of glutathione could directly sensitize to GPX4 inhibition. Alternatively, diverse UPR stressors may upregulate one or more specific proteins that enhance ferroptosis sensitivity indirectly. While ATF4 appeared to oppose GPX4 inhibitor sensitization, other proteins could promote this effect. Of note, the lipid metabolic protein membrane-bound O-acyltransferase domain-containing 7 (MBOAT7) was one of fifty proteins that increased from undetectable basal levels in both HT-1080^N^ and U-2 OS^N^ cells treated with carfilzomib. MBOAT7 can promote ferroptosis sensitivity by altering polyunsaturated fatty acid (PUFA)-containing phospholipids levels.^86^ AlteredPUFA metabolism downstream of proteasome inhibition could account for heightened lipid peroxidation and ferroptosis following GPX4 inhibitor treatment, and this will be examined in future studies. Regardless, our results suggest that combining clinical proteasome inhibitors like carfilzomib and emerging GPX4 inhibitors like compound 28 (ref.^50^) for cancer therapy could be an intriguing translational possibility.

### Limitations of the study

We are not the first to suggest that the proteasome may regulate ferroptosis sensitivity in some way^8,17,20,87,88^, but the literature in this area is contradictory and has been difficult to parse. Our findings suggest at least three possible explanations for existing discrepancies. First, cell type-specific regulation may render proteasome function differentially important for ferroptosis. Our cell models generally all yielded similar phenotypes, but we have only examined established cancer cell lines. It is possible that freshly isolated cancer cells or normal primary cells^20^ respond differently to proteasome inhibition. Second, we show here in our models that proteasome inhibition can sensitize to direct inhibition of GPX4 yet at the same time produces resistance to system x_c_ ^-^ inhibition. It is important to carefully consider that responses to different ferroptosis inducers can vary substantially.^30,40^ Third, studying how proteasome perturbation impacts ferroptosis is made difficult by the fact that proteasome inhibition causes substantial confounding baseline apoptosis. Inhibiting the execution of apoptosis execution helps surmount this technical challenge but could also rewire cell signaling in unexpected ways we have not examined here.

## ACKNOWLEDGEMENTS

We thank Jaclyn Ng, Cassandra Stawicki, Arianna Silva-Torres, Celeste Riepe, Magda Wąchalska, Jan Carette and Monther Abu-Remaileh for insightful discussions and/or reagents, Jiangbin Ye for sh-NT and sh-*ATF4* cell lines, Michael Lee for U-2 OS wild-type and BAX/BAK1 double knockout cell lines, Joshua Arribere for the proteasome reporter plasmid, Gina DeNicola for reagents to examine CHAC1, and Livnat Jerby for cells. We thank A. Joseph, W. Lee, N. Haseley, P. Sutton, and I. Srivastava for comments on the manuscript. This work was supported by awards from the Grace Science Foundation, and from the National Institutes of Health to M.B.M. (F31CA284784), O.B. (R35GM153301), and S.J.D. (R01GM122923).

## AUTHOR CONTRIBUTIONS

M.B.M. and S.J.D. conceived and wrote the manuscript. M.B.M., R.U., K.S., A.G., and J.G. performed experiments. G.C.F., A.V., and O.B. contributed reagents.

M.B.M. and S.J.D. wrote the manuscript with input from all co-authors.

## METHODS

### RESOURCE AVAILABILITY

#### Lead Contact

Further information and requests for resources and reagents should be directed to and will be fulfilled by the Lead Contact, Scott Dixon.

#### Materials Availability

Plasmids, cell lines and other materials generated in this study will be shared by the lead contact upon request.

#### Data and Code Availability

- Uncropped immunoblots will be made available online at the Mendeley Data Repository at the date of publication.
- Any additional information required to reanalyze the data reported in this paper is available from the lead contact upon request.

### METHODS DETAILS

#### Cell lines

HT-1080^N^ (sex: male), NCI-H1299^N^ (sex: male), U-2 OS^N^ (sex: female), and T98G^N^ (sex: male) were described previously.^43,64^ HT-1080^N^ Control and *GPX4*^KO1/2^ cell lines were also previously described.^51^ HT-1080^N^ Control and *NFE2L1*^KO1/2^ cell lines were developed as described below. Lenti-X and NCI-H1299 cells were kind gifts from Livnat Jerby (Stanford School of Medicine) and Gina DeNicola (Moffitt Cancer Center), respectively. HT-1080 cells were cultured in medium containing DMEM (Cat# 10-013-CV, Corning), 10% FBS (Cat# A5670701, Thermo Fisher Scientific), 1% penicillin/streptomycin (Cat# 15-070-063, Fisher Scientific), and 1% non-essential amino acids (Cat# 11140050, Thermo Fisher Scientific). U-2 OS cells were cultured in McCoy’s 5A medium (Cat# 10-050-CV, Corning), 10% FBS, and 1% penicillin/streptomycin. Lenti-X, H1299, and T98G cells were cultured in DMEM, 10% FBS, and 1% penicillin/streptomycin.

#### Chemicals and reagents

ML162 was synthesized by Acme (Palo Alto, CA). Compound 28 (ref.^50^) was synthesized and validated using mass spectrometry and proton nuclear magnetic resonance by Pharmaron (Beijing, China). Erastin2 (Cat# HY-139087), ONX-0914 (Cat# HY-13207), thapsigargin (Cat# HY-13433), epoxomicin (Cat# HY-13821), Q-VD-OPh (Cat# HY-12305), and tegavivint (Tegatrabetan, Cat# HY-109103) were purchased from MedChemExpress. M-3258 (Cat# 38160), bortezomib (Cat# 10008822), vacquinol-1 (Cat# 16321), and edelfosine (Cat# 60912) were obtained from Cayman Chemical Company. Carfilzomib (Cat# S2853), 1*S*,3*R*-RSL3 (Cat# S8155), camptothecin (Cat# S1288), JTC-801 (Cat# S2722), and zinc pyrithione (Cat# S4075) were purchased from Selleck Chemicals (Houston, TX). Cycloheximide (CHX, Cat# 01810), urea (Cat# U5378-500G), tunicamycin (Cat# T7765), and N-acetyl-L-cysteine (Cat# A8199) were obtained from Millipore Sigma. Ferrostatin-1 (Cat# SML0582) was obtained from Sigma-Aldrich. Emetine dihydrochloride was purchased from Hello Bio (Cat# HB7389). TPEN was a kind gift from Monther Abu-Remaileh.

#### Genome-wide CRISPR-Cas9 screens

Genome-wide pooled CRISPR/Cas9 screens were conducted using the ChemoGenix platform (IRIC, Université de Montréal; https://chemogenix.iric.ca/) as previously described.^31^ Briefly, a NALM-6 clone bearing an integrated inducible Cas9 expression cassette was transduced with the genome-wide EKO sgRNA library which contains 278,754 different individual sgRNAs.^31^ Cell stocks were made and frozen in LN2. For each screen, an aliquot of frozen cells was thawed in 10% FBS RPMI for 1 d, then the addition of 2 µg/mL doxycycline for 7 d initiated Cas9 expression. The population was then split into different T-75 flasks in a volume of 70 mL at a density of 4 × 10^5^ cells/mL. A total of 28 × 10^6^ cells were used per flask/screen, equivalent to 100 cells/sgRNA. Cells were treated with DMSO (vehicle control), RSL3 or erastin2 (from 1,000x stock solutions dissolved in DMSO) for 8 d. Cells were counted every other day and diluted back to 4 × 10^5^ cells/mL, with the addition of more compound, whenever cells reached 8 × 10^5^ cells/mL. At the end of the experiment, cells were harvested by centrifugation and the Gentra Puregene kit (QIAGEN, Germantown, MD) was used to extract genomic DNA according to manufacturer’s instructions. PCR was used to amplify sgRNA sequences as described.^31^ Next generation sequencing (Illumina NextSeq 2000) was used to determine sgRNA frequencies. Total read counts per sgRNA were obtained from sequence reads aligned using Bowtie2.2.5 in the forward direction only (–norc option), with default parameters otherwise. A modified version of the RANKS algorithm^31^, or Context-dependent Robust Analytics and Normalization for Knockout Screens (CRANKS), was used to compute chemogenomic interaction scores.

#### Analysis of *PSMB8* and *PSMB5* gene expression across cancer cell lines

Gene expression data for *PSMB8* and *PSMB5* in catalogued cancer cell lines was downloaded from the DepMap portal (https://depmap.org/portal/) on August 20, 2025, and plotted using GraphPad Prism.

#### Cell death measurement

For experiments performed in 384-well plates, cells were seeded on day 1 at a concentration of 1,500 cells/well (1,000 cells/well for NCI-H1299 cells) for experiments where cell death would be observed starting on day 2 until day 4 (48 h), and at a density of 750 cells/well for experiments where cell death would be observed starting on day 2 until day 5 (72 h). For experiments performed in 96-well plates, cells were typically seeded on day 1 at a density of 8,000 cells per well. Assay plates were briefly centrifuged after seeding to settle cells in the bottom of the wells. For most experiments, cells were treated with compounds on day 2 in medium containing SYTOX Green (20 nM, Cat# S7020, Thermo Fisher Scientific) and plates were briefly centrifuged. For the experiment in which HT-1080^N^ cells were treated with carfilzomib in addition to a panel of lethal compounds (including vacquinol-1, JTC-801, etc.), cells were seeded on day 1 and treated with the indicated compounds on day 3 prior to imaging. For the HT-1080^N^ Control and GPX4^KO^ ferrostatin-1 titration assay, cells were seeded as described above. Prior to treatment with lethal compounds, cell death inhibitors, and SYTOX Green, each plate underwent three rounds of medium removal followed by washing with 1X PBS.

An Incucyte S3 live cell analysis instrument (Sartorius, Göttingen, Germany) housed within a tissue culture incubator maintained at 37°C with 5% CO2 was used to image wells (800 ms acquisition time for red for cells expressing mKate2, 400 ms acquisition time for green, 10X objective) every 6-24 h for 48-72 h total. For non-tskMP HT-1080^N^ and U-2 OS^N^ experiments, the following parameters were used: green (segmentation: adaptive, threshold adjustment: 2-6 GCU, edge split: on, edge sensitivity: 10, hole fill: 10 µm^2^, minimum area: 40 µm^2^, maximum area: 2500 µm^2^), red (segmentation: adaptive, threshold adjustment: 1-2 RCU, edge split: on, edge sensitivity: -30, hole fill: 2 µm^2^, minimum area: 20-40 µm^2^, maximum area: 3000 µm^2^, maximum eccentricity: 1, minimum mean intensity: 6), and yellow (minimum area: 70-100 µm^2^, maximum area: 5000 µm^2^, minimum green mean intensity: 3-10, minimum red mean intensity: 3-10). For U-2 OS cells, the following parameters were used: green (segmentation: adaptive, threshold adjustment: 6 GCU, edge split: on, edge sensitivity: 10, hole fill: 10 µm^2^, minimum area: 40 µm^2^, maximum area: 2500 µm^2^) and phase (segmentation: AI confluence, hole fill: 20 µm^2^, minimum area: 100 µm^2^). For H1299^N^ cells, the following parameters were used: green (segmentation: adaptive, threshold: 4 GCU, edge split: on, edge sensitivity: -5, hole fill: 20 µm^2^, minimum area: 30 µm^2^, maximum area: 2500 µm^2^), red (segmentation: adaptive, threshold adjustment: 7 RCU, edge split: on, edge sensitivity: -35, hole fill: 100 µm^2^, minimum area: 40 µm^2^, maximum area: 3000 µm^2^, minimum mean intensity: 6), and yellow (minimum area: 100 µm^2^, maximum area: 5000 µm^2^, minimum green mean intensity: 10, minimum red mean intensity: 10). For H1299 cells, the following parameters were used: green (segmentation: adaptive, threshold adjustment: 9 GCU, edge split: on, edge sensitivity: -5, hole fill: 20 µm^2^, minimum area: 100 µm^2^, maximum area: 2500 µm^2^) and phase (segmentation: AI confluence). For the experiment in which both H1299^N^ and T98G^N^ cells were analyzed, the following parameters were used: green (segmentation: adaptive, threshold adjustment: 2 GCU, edge split: on, edge sensitivity: 10, hole fill: 10 µm^2^, minimum area: 40 µm^2^, maximum area: 2500 µm^2^), red (segmentation: adaptive, threshold adjustment: 1 RCU, edge split: on, edge sensitivity: -30, hole fill: 2 µm^2^, minimum area: 20 µm^2^, maximum area: 3000 µm^2^), and yellow (minimum area: 70 µm^2^, maximum area: 5000 µm^2^, minimum green mean intensity: 3, minimum red mean intensity: 3). The lethal fraction was calculated as described^89^. For cells without mKate2 expression, cell death was quantified by calculating the maximum SYTOX Green signal over the percent confluence. For U-2 OS control and *BAX BAK1* double KO cells, max(SYTOX Green) was calculated over 0 and 48 h timepoints. For HT-1080 and H1299 cells, max(SYTOX Green) was calculated over 0, 24, and 48 h timepoints. To calculate Bliss deviation scores from max(SYTOX Green)/Confluence-based data, AUC values were normalized such that each value was between 0 and 1.

#### Cystine deprivation and supplementation

Experiments were conducted by replacing growth medium with starvation medium, supplemented with the relevant amino acid stock solutions. DMEM minus cystine was constituted by supplementing DMEM for SILAC (Cat# 21013024, Thermo Fisher Scientific), lacking L-glutamine, L-methionine and L-cystine, with a 100X stock solution of L-glutamine (Cat# G3126, Sigma-Aldrich) at 52 g/L and a 1000X stock solution of L-methionine (Cat# M9625, Sigma-Aldrich) at 26.7 g/L. To generate ‘complete medium’, a 1000X stock solution of L-Cystine (Cat# C8755, Sigma-Aldrich) at 56 g/L was also added. To minimize the contribution of monomeric amino acids contained in normal FBS, all media were supplemented with 10% dialyzed FBS (dFBS, Cat# 26400044, Thermo Fisher Scientific) and 1% P/S.

#### Lentivirus production

The following protocol was used to generate lentivirus. Lentivirus was used to package the following plasmids: pLenti CMV Puro DEST empty vector (‘pLenti CMV-EV’, kindly provided by Jan Carette, Cat# 17452, Addgene) or pLenti CMV-MCL1 (described previously)^90^, or a pLenti eIF1α mCherry-T2A-eGFP plasmid (‘pLenti mCherry-T2A-eGFP’, the kind gift of J. Arribere, University of California, Santa Cruz)^52^. On day 1, 6 cm dishes were incubated with 1X poly-D-lysine (Cat# A3890401) for 20 min at room temperature and subsequently washed twice with 1X PBS. Lenti-X cells were seeded into these dishes in complete growth medium such that they reached approximately 80% confluence after one day. On day 2, the complete growth medium was replaced with medium lacking P/S. A solution consisting of 2 µg of the pLenti

CMV-EV or pLenti CMV-MCL1 in addition to 1.2 µg psPAX2 (Cat# 12260, Addgene), 600 ng pMD2.G (Cat# 12259, Addgene), and 12 µL TransIT-Lenti Transfection Reagent (Cat# MIR 6604, Mirus Bio) in a total volume of 400 µL Opti-MEM (Cat# 31985070, Life Technologies) was incubated for 10 min at room temperature then added drop-wise to the Lenti-X cells. 6 h post-transfection, the existing medium was removed and 4 mL fresh medium (without P/S) containing 8 µL of the Viralboost Reagent (Cat# VB100, Alstem Cell Advancements) was added to each 6 cm dish. On day 4, viral supernatants were collected and centrifuged for 5 min at 450 *x g*. The supernatants were then passed through a 0.45 µm PVDF filter (Cat# 09-720-4, Fisher Scientific) prior to a fresh tube prior to aliquoting and storage at -80°C.

#### Lentiviral transduction

Lentivirus was used to deliver Incucyte Nuclight R Lenti (bleo) (Cat# 4478, Sartorius), pLenti CMV-EV or pLenti CMV-MCL1, and pLenti mCherry-T2A-eGFP to cells. To transduce cells, 20,000 HT-1080 (for Incucyte Nuclight R or pLenti mCherry-T2A-eGFP transduction) HT-1080^N^ (for pLenti CMV-EV or pLenti CMV-MCL1 transduction) cells were seeded per well of a 12-well plate. The following day, cells were transduced with lentivirus and 8 µg/mL hexadimethrine bromide (aka polybrene). The next day, the existing medium was replaced with fresh medium. The day after, successfully transduced cells were isolated using fluorescence-activated cell sorting (for cells transduced with Incucyte Nuclight R) or 1.25 µg/mL puromycin (for cells transduced with pLenti CMV-EV, pLenti CMV-MCL1, or pLenti mCherry-T2A-eGFP).

#### Generation of monoclonal knockout line

Single guide RNAs (sgRNA) directed against exon 2 were used to generate the HT-1080^N^ *NFE2L1* gene-disrupted cell lines; forward, 5′-CACCgTTTCTCGCACCCCGTTGTCT-3′ and reverse, 5′-AAACAGACAACGGGGTGCGAGAAAc-3′. sgRNAs were cloned into pSpCas9(BB)-2A-GFP, a kind gift from Jan Carette, Stanford University School of Medicine, as described.^51^ Briefly, primers were annealed by heating to 95 °C, then cooled by 2.5 °C/min to 25 °C. The resulting oligo duplex was diluted 1:200 in ddH2O and used for cloning into the pSpCas9(BB) plasmid. The ligation reaction contained the following: 100 ng pSpCas9(BB), 2 μL of the diluted oligo duplex, 2 μL 10× FastDigest buffer (Thermo Fisher Scientific), 1 μL 10 mM dithiothreitol, 1 μL 10 mM adenosine triphosphate, 1 μL FastDigest BpiI (D1014; Thermo Fisher Scientific), 0.5 μL T4 DNA ligase (EL0014; Thermo Fisher Scientific), and 12.5 µL ddH2O to produce a final reaction volume of 20 µL. The ligation reaction was incubated for 1 h ([37°C, 5 min; 21°C, 5 min] × six cycles). Two microliters of the reaction product were used to transform competent DH5α cells on LB-amp (100 µg/mL) plates. Plasmid DNA was extracted using a spin column (27106; Qiagen) and confirmed via Sanger sequencing. Cells were transfected with 2.5 µg DNA using Opti-MEM I Reduced Serum Medium (31985-062; Life Technologies) and Lipofectamine LTX Reagent with PLUS Reagent (15338-100; Life Technologies), according to the manufacturer’s instructions. The next day the medium was replaced, and the cells were allowed to recover for 24 h. Transfected cells were then trypsinized, washed in 1× PBS, and sorted for green fluorescent protein–positive (GFP+) cells with a Carmen cell sorter (BD Biosciences) (Stanford Shared FACS Facility) into a 96-well plate prefilled with DMEM containing 30% FBS and 0.5 U/mL P/S. Cells were incubated at 37°C until distinct clones appeared. Individual clones were amplified in normal (10% FBS) medium to produce enough cells for protein analysis. Clones were validated by Western blot to confirm loss of protein expression.

#### Generation of polyclonal knockout lines

On day 1, 600,000 H1299 Cas9 cells were seeded in a 10 cm dish. On day 3, cells were nucleofected with 400 ng of pLenti-CRISPR-V2 plasmids harboring Cas9 as well as sg*Control* or sg*CHAC1* (kindly gifted by Gina DeNicola) using the SF Cell Line 4D-Nucleofector X Kit S kit (Cat# V4XC-2032, Lonza) according to manufacturer’s instructions. On day 5, successfully nucleofected cells were selected using 1 µg/mL puromycin.

#### siRNA transfection

*PSMB5* siRNAs (Cat# SASI_Hs01_00076893 and Cat# SASI_Hs01_00076894) and *CYCS* siRNAs (Cat# SASI_Hs01_00190690 and Cat# SASI_Hs01_00190691) were purchased from Millipore Sigma. AllStars Negative Control siRNA (‘siNeg’, Cat# 1027280, Qiagen) was used as a control in experiments involving siRNA. On Day 1, HT-1080^N^ cells were reverse transfected with siRNA in either 6-well plates or 6 cm dishes. Briefly, per well of a 6-well plate, 500 µL OptiMEM (Cat# 31985-070, Thermo Fisher Scientific), 2.5 µL Lipofectamine RNAiMAX (Cat# 13778-075, Thermo Fisher Scientific), and 20 pmol siRNA were used. For 6 cm dishes, these volumes were doubled. For si*PSMB5* experiments, cells destined for the cell death assay were treated with DMSO (vehicle control) or 25 µM Q-VD-OPh at the time of cell seeding. For 6-well plates, cells were seeded at a density of 75,000 cells per well in a total of 2 mL volume and for 6 cm dishes at a density of 150,000 cells per dish in a total of 5 mL volume. On Day 3, cells destined for the cell death assay were harvested via trypsinization and 62,000 cells were seeded in each well of a 12-well plate. For si*PSMB5* experiments, cells were again treated with either DMSO (vehicle control) or 25 µM Q-VD-OPh. For cells destined for analysis by western blotting, on Day 3, cells were harvested via trypsinization (500 µL for 6 cm dishes) and processed as per usual (see western blotting section). For the cell death experiments, on Day 4, medium was aspirated from the 12-well assay plates and cells were treated with the indicated compounds. Assay plates were imaged every 12 h to 24 h for a total of 48 h using an S3 live cell analysis instrument with a 10X objective. Red, green, and yellow objects were quantified and lethal fraction was calculated as detailed in the cell death measurement section.

#### *GPX4*^KO^ cell death analysis

For routine culture, HT-1080^N^ Control, *GPX4*^KO1^ and *GPX4*^KO2^ cell lines were grown in medium containing 1 µM ferrostatin-1. For experiments, on Day 1, cells were seeded in 96-well plates at a density of 8,000 cells/well in 200 µL medium (Corning, Cat# 3598) containing 1 µM ferrostatin-1. The assay plates were briefly centrifuged to collect cells in the bottom of the wells. On Day 2, cells were washed three times with 1X PBS. After the third wash, PBS was removed and medium containing SYTOX Green and either DMSO (vehicle control) or Q-VD-OPh was added to the cells. Next, medium containing DMSO (vehicle control) or carfilzomib was then added to the cells. Finally, medium containing DMSO (vehicle control) or ferrostatin-1 was added to the cells. The final concentrations of each compound in the assay plates were: 20 nM SYTOX Green, 25 µM Q-VD-OPh, 10 nM carfilzomib, and eleven-point, 4-fold dose-points of ferrostatin-1 starting at a high dose of 1 µM. The assay plates were then briefly centrifuged prior to imaging every 6 h for a total of 48 h at 10X magnification using the Sartorius S3 live cell analysis instrument. For analysis, live cells (red objects), dead cells (green objects), and double positive (yellow objects) were counted and the lethal fraction was quantified as described above.

#### Time-staggered kinetic modulatory profiling (tskMP)

On day 1, HT-1080^N^ and U-2 OS^N^ cells were seeded at a density of 750 cells per well of a 384-well plate and briefly centrifuged to settle cells in the bottom of the wells. On day 2, the 24 h pretreatment wells were treated with carfilzomib at final concentrations of 1 nM, 10 nM, or 100 nM and the plates were centrifuged briefly. On day 3, the 4 h, 1 h, and 0 h pretreatment wells were treated with the above concentrations of carfilzomib. At time “0” (i.e., the end of the pretreatments), The medium was replaced with medium containing DMSO (vehicle control) or 10-point, 2-fold (or 4-fold for some experiments, as noted) dilution series of erastin2 (high dose 5 µM) or RSL3 (high dose 5 µM). Medium also contained SYTOX Green (20 nM) and DMSO (vehicle control), ferrostatin-1 (1 µM), or Q-VD-OPh (25 µM), with final concentrations listed in brackets. The assay plates were briefly centrifuged and then imaged using a Sartorius S3 instrument every 6 h for a total of 48 h. Images were acquired using the 10X objective with the following exposures: red channel (800 ms) and green channel (400 ms). For HT-1080^N^ cells, the following image analysis parameters were used: green (segmentation: adaptive, threshold adjustment: 6 GCU, edge split: on, edge sensitivity: 10, hold fill: 10 µm^2^, minimum area: 40 µm^2^, maximum area: 2,500 µm^2^), red (segmentation: adaptive, threshold adjustment: 2 RCU, edge split: on, edge sensitivity: -30, hole fill: 2 µm^2^, minimum area: 40 µm^2^, maximum area: 3,000 µm^2^, maximum eccentricity: 1, minimum mean intensity: 6), yellow (minimum area: 100 µm^2^, maximum area: 5,000 µm^2^, minimum green mean intensity: 10, minimum red mean intensity: 10). For U-2 OS^N^ cells, the following image analysis parameters were used: green (segmentation: adaptive, threshold adjustment: 6 GCU, edge split: on, edge sensitivity: -5, hole fill: 10 µm^2^, minimum area: 40 µm^2^, maximum area: 2500 µm^2^), red (segmentation: adaptive, threshold adjustment: 2 RCU, edge split: on, edge sensitivity: -30, hole fill: 2 µm^2^, minimum area: 40 µm^2^, maximum area: 3,000 µm^2^, maximum eccentricity: 1, minimum mean intensity: 6), yellow (minimum area: 100 µm^2^, maximum area: 5,000 µm^2^, minimum green mean intensity: 10, minimum red mean intensity: 10). Lethal fraction was calculated as described in the ‘cell death measurement’ section.

#### Western blotting

For the HT-1080 shNT and sh*ATF4* western blots, cells were seeded at a density of 5.3 × 10^5^ cells per 6 cm dish on Day 1. For the HT-1080^N^ control and *NFE2L1*^KO^ western blots, cells were seeded at a density of 275,000 cells per well of a 6-well plate on Day 1. For the western blot experiment in which HT-1080^N^ cells were treated with combinations of RSL3, carfilzomib, and cycloheximide, cells were seeded at a density of 1.4 × 10^6^ cells per 10 cm dish on Day 1. For H1299 sg*CHAC1* validation, 1.8 × 10^5^ cells were seeded per well of a 6-well plate. On Day 2, cells were treated with the indicated compounds for the indicated treatment duration.

Cells were then harvested using the following methods. Cells were washed with 1X PBS then incubated in trypsin for 4 min. Trypsin was quenched with growth medium and cells were collected in 1.5, 2 mL, or conical tubes, as appropriate. Samples were centrifuged at 350-750 *x g* for 4 min, then washed with 1X PBS prior to centrifugation at 350-750 *x g* for 4 min. PBS was then aspirated from each sample and 9M urea (Millipore Sigma, Cat# U5378) was added prior to freezing at -20C until needed. Thawed lysates were sonicated ten times for 1 s each at 60% amplitude with a Fisher Scientific Model 120 sonic dismembrator (Thermo Fisher Scientific). Debris was pelleted by centrifugation at 18,213 *x g* for 15 min at 4°C. Supernatants were then transferred to fresh 1.5 mL tubes. To quantify the concentration of protein in each sample, a BCA assay (Thermo Fisher Scientific, Cat# 23224 and Cat# 23228) was performed in a 96-well plate (Corning, Cat# 07-200-103 or Cat# 07-200-760), according to the manufacturer’s instructions. A Synergy Neo2 multimode plate reader (BioTek Instruments, Winooski, VT) was used to measure each samples’ absorbance at 562 nm.

For cells that did not require treatment, cells were harvested during cell passaging. For HT-1080^N^ control and *MCL1* overexpression lines, 300,000 cells were collected, for HT-1080^N^ Control and *GPX4*^KO^ and U-2 OS Control and *BAX*/*BAK1* DKOs, 500,000 cells were collected. Cells were then processed as described above.

To prepare samples for electrophoresis, 4X Bolt LDS Sample Buffer (Thermo Fisher Scientific, Cat# B0007) and 10X Bolt Sample Reducing Agent (Thermo Fisher Scientific, Cat# B0009) were added to each sample. On day 1 of western blotting, samples were boiled for 5 min at 95°C. Samples were then run alongside the Chameleon Duo Pre-stained Protein Ladder (Cat# NC0738562, Fisher Scientific) using Bolt 4-12% Bis-Tris Plus Gels (Cat# NW04120BOX or Cat# NW04122BOX or Cat# NW04125BOX, Thermo Fisher Scientific) and 1X Bolt MES SDS Running Buffer (Cat# B0002, Life Technologies) for 1 h 45 min at 100V. Either the iBlot2 or iBlot3 system (Life Technologies) was used to transfer proteins from the gel to a nitrocellulose membrane. A 0.5 h incubation in Intercept (TBS) blocking buffer (Cat# 927-6001, LICORbio, Lincoln, NE) was used to block the blots. Blots were then incubated in the respective primary antibodies, diluted in blocking buffer, overnight at 4°C. The following primary antibodies were used: rabbit anti-ATF4 (1:1,000, Cat# 11815, Cell Signaling Technology, Danvers, MA), rabbit anti-BAX (1:1,000, Cat# 2772, Cell Signaling Technology, Danvers, MA), rabbit anti-BAK (1:1,000, Cat# 3814, Cell Signaling Technology, Danvers, MA), mouse anti-GAPDH (1:10,000, Cat# 60004-1-Ig, Proteintech, Rosemont, IL), rabbit anti-CYCS (1:1,000, Cat# 11940, Cell Signaling Technology, Danvers, MA), rabbit anti-GPX4 (1:1,000, EPNCIR144, Cat# 125066, Abcam), rabbit anti-PSMB5 (Cat# 12919, Cell Signaling Technology, Danvers, MA), rabbit anti-DDIT3/CHOP (1:1,000, Cat# 15204-1-AP, Proteintech, Rosemont, IL), and rabbit anti-CHAC1 (1:500, Cat# 15207-1-AP, Proteintech, Rosemont, IL). On day 2 of western blotting, Tris-buffered saline (TBS, Cat# 0788, ISC BioExpress, Kaysville, UT) with 0.1% Tween 20 (TBST) was used to wash the blots. Blots were washed three times for 4 min each. Blots were then incubated in a 1:15,000 dilution of secondary antibodies in Intercept (TBS) blocking buffer) for 1 h at room temperature. Secondary antibodies included: 680LT Donkey anti-mouse (Cat# 926-68022, LICORbio) and 800CW Donkey anti-rabbit (Cat# 926-32213, LICORbio). Membranes were then washed three times for 4 min each with TBST at room temperature. Blots were imaged with a LICORbio Odyssey M machine.

#### Puromycylation assay

On day 1, HT-1080^N^ cells were seeded at a density of 460,000 cells per 6 cm dish. On day 2, cells were treated with regular medium (no puromycin control) or DMSO or 100 nM carfilzomib plus DMSO or 35 µM cycloheximide for 10 h. 15 min prior to harvesting, cells were treated with a final concentration of 5 µg/mL puromycin. To harvest, medium was aspirated from the cells, cells were washed with cold 1X PBS, and PBS was aspirated from the cells. Cells were then incubated in 250 µL complete RIPA buffer including 1:100 protease inhibitor cocktail (Millipore Sigma, Cat# P8340) and NaF (Millipore Sigma, Cat# S6776) for 20 min. Next, cells were scraped and collected in 1.5 mL tubes and centrifuged at 13,000 *x g* for 20 min at 4C. Supernatants were then transferred to fresh 1.5 mL tubes and stored at -20C short-term before western blotting. Overall, protein quantification, protein separation via polyacrylamide gel, and transfer steps were completed as described above with the following exception: 20 µg total protein was loaded into each well of the polyacrylamide gel. After transferring proteins to nitrocellulose, blots were washed three times in HPLC-grade water for 1 min each prior to a 15 min incubation in Ponceau S Staining Solution. The blots were then washed for 1 min with HPLC-grade water and imaged using a LICORbio Odyssey M machine. After imaging, blots were washed three times with TBS-T for 10 min each prior to a 1 h incubation in blocking buffer. Blots were then washed three times with TBS-T for 4 min each, then probed with an anti-puromycin antibody (Cat# MABE343, Millipore Sigma) at a dilution of 1:10,000 in blocking buffer overnight at 4C. The next day, blots were washed three times with TBS-T for 4 min each, incubated in a donkey anti-mouse secondary antibody (Cat# 926-32212, LICORbio) at a dilution of 1:15,000 in blocking buffer for 1 h, and again washed three times with TBS-T for 4 min each. Blots were stored in 1X PBS and imaged using the LICORbio Odyssey M machine. To probe for GAPDH, blots were stripped with 1X nitrocellulose stripping buffer (Cat# 928-40030, LICORbio, diluted with MilliQ water) for 15 min. Blots were then washed briefly six times with 1X PBS, incubated in blocking buffer for 30 min, and probed with 1:1,000 GAPDH antibody (Cat# 2118, Cell Signaling Technology) in blocking buffer overnight at 4C. The following day, blots were washed three times with TBS-T for 4 min each, incubated in a 1:15,000 dilution of donkey anti-rabbit secondary antibody (Cat# 926-32213, LICORbio) in blocking buffer for 1 h, washed three times with TBS-T for 4 min each, stored in 1X PBS, and imaged using a LICORbio Odyssey M machine. All blot images were quantified as described above.

#### Proteomic analysis

On day 1, HT-1080^N^ and U-2 OS^N^ cells were seeded in duplicate wells of a 6-well plate. On day 2, cells were treated with DMSO or 100 nM carfilzomib for 15 h (for HT-1080^N^) or 22 h (for U-2 OS^N^), which accorded to approximately 0.5 h prior to death onset, respectively. On day 3, cells were harvested as follows. After two washes with 1 mL cold 1X PBS each, 500 µL cold 1X PBS was added to each well and cells were lifted using a cell scraper. The cell suspension was then added to a 1.5 mL tube, cells were pelleted by centrifugation at 500 x *g* for 5 min at 4°C, PBS was aspirated from the cell pellet, and samples were frozen at -80°C.

Samples were then analyzed by mass spectrometry at the University of California Davis proteomics core facility. At the facility, cells were lysed in a solution containing 5% SDS and 50 mM triethyl ammonium bicarbonate and with bead beating. A BCA assay (Thermo Fisher Scientific, Cat# 23225) was used to quantify protein in each sample. Samples underwent clean-up, reduction, alkylation, and tryptic proteolysis with suspension-trap (ProtiFi) devices, a form of filter-aided sample preparation in which acid-aggregated proteins are trapped in a quartz filter before enzymatic proteolysis. To resuspend the proteins, 50 µL of buffer, containing 5% SDS and 50 mM TEAB (pH 7.55), was added to each sample. Samples were treated with dithiothreitol to reduce disulfides, and subsequently alkylated, protected from light, using 50 mM TEAB buffer containing iodoacetamide. Samples were digested for 4 h at 37°C with trypsin at a ratio of 1:100 enzyme:protein (wt/wt). This reaction was terminated by adding 1% trifluoroacetic acid. The eluted tryptic peptides were dried using a vacuum centrifuge and the peptides reconstituted in a solution of water and 2% acetonitrile (ACN). Equal amounts of peptide were used for each sample in subsequent analysis. First, peptides were loaded onto a disposable Evotip C18 trap column (Evosep Biosystems, Denmark) according to the manufacturer’s directions. 2-proponal was used to wet the Evotips that were subsequently equilibrated with 0.1% formic acid. Evotips were then loaded by centrifugation at 1,200 x *g*. Next, 0.1% formic acid was used to wash the Evotips prior to the addition of 200 µL 0.1% formic acid to each Evotip to maintain moisture. An Evosep One instrument (Evosep Biosystems) was used to perform nanoLC on each of the loaded samples. Tips were eluted directly onto a PepSep analytical column (dimensions: 150 µm x 25 cm, stationary phase: C18, pore size: 1.5 µm/100 Å, Bruker Daltonics, Bremen, Germany) and a ZDV spray emitter (Bruker Daltonics). Mobile phase A consisted of 0.1% formic acid in water (v/v). Mobile phase B was comprised of 80/20/0.1% CAN/water/formic acid (v/v/v). Each run was 21 min as part of a 60 samples-per-day standard pre-set approach.

Mass spectrometry was conducted using a hybrid trapped ion mobility spectrometry-quadrupole time of flight mass spectrometer (timsTOF HT, Bruker Daltonics) in PASEF mode. The timsTOF machine was modified with a nano-electrospray ion source (CaptiveSpray, Bruker Daltonics). Desolvated ions traveled through a glass capillary to the vacuum region and were deflected into the TIMS tunnel, consisting of two parts (dual TIMS) separated electrically. The first part was used as an ion accumulation trap that primarily stored all ions entering the mass spectrometer. The second part performed trapped ion mobility analysis. The dual TIMS analyzer was operated at a fixed duty cycle close to 100% using equal accumulation and a ramp time of 85 ms. To perform data-independent analysis (DIA), an MS scan was performed. Subsequently, MSMS scans were performed with 36 precursor windows at a width of 25^Th^ per 1.09 s cycle, over the mass range of 300-1200 Dalton. The TIMS scans layer the doubly and triply charged peptides over an ion mobility -1/k0- range of 0.7-1.3 V*s/cm^2^. The collision energy was ramped linearly as a function of the mobility from 59 eV at 1/K0=1.4 to 20 eV at 1/K0=0.6. Spectronaut 18.4 (Biognosys Schlieren, Switzerland) direct DIA workflow with PTM localization was used to analyze the DIA data. The enzyme parameter was set to trypsin/P Specific and the method allowed for two missed cleavages. For Carbamidomethyl, fixed modifications were used. For Acetyl (Protein N-term) and Oxidation, variable modifications were used. PSM and Protein Group FDR were set to 0.01% to perform DIA search identification. To quantify a given protein, at least two peptides were required per protein group.

Analysis of proteins detected upon carfilzomib treatment was performed as follows. For HT-1080^N^ and U-2 OS^N^ cells, proteins only detected upon carfilzomib treatment were plotted in order of ascending abundance. For functional interaction analysis, proteins only detected in carfilzomib-treated HT-1080^N^ and U-2 OS^N^ cells were entered into the STRING database string-db.org). The proteomics species labeled “UPK3BL1;UPK3BL2” was split into “UPK3BL1” and “UPK3BL2” for analysis. The same approach was used for “FAM72C;FAM72D” and “POLR2J2;POLR2J3”. MSANTD7 and HSPA7 were not present in the STRING database and omitted from further analysis. Default settings were used for the analysis (merge rows by term similarity: Don’t merge, maximum FDR shown: FDR <= 0.05, maximum signal shown: Signal >= 0.01, Maximum strength shown: Strength >= 0.01, minimum count in network: 2, statistical background: Whole genome). The protein interaction map was exported to Cytoscape for adjustments. Proteins that had no interactors were omitted from the map. FAM72C and FAM72D were omitted, as they only interacted with one another (and were originally detected as “FAM72C;FAM72D” in the proteomics analysis). POLR2J2 and POLR2J3 were omitted for the same reasons. Node size was normalized to the average abundance of each protein across HT-1080^N^ and U-2 OS^N^ cells upon carfilzomib treatment. Specifically, average abundance was log transformed and all proteins were normalized to the average logged abundance of the least abundant protein (CEP68). These normalized values were multiplied by 37 to generate the node sizes for each protein. Node color was scaled by the number of interactions (darker = more interactions).

#### Glutathione measurements

Total glutathione (GSSG + GSH) abundance was measured using the Glutathione Assay Kit (Cat# 703002) from Cayman Chemical Company. On day 1, HT-1080^N^ cells were seeded at a density of 160,000 cells per well of a 6-well plate. On day 2, cells were treated with 300 nM erastin2 or DMSO or 100 nM carfilzomib +/- 35 µM cycloheximide for 10 h. Immediately prior to harvest, cells were imaged using an S3 live cell analysis instrument (red channel acquisition time: 800 ms, green channel acquisition time: 400 ms, 10X objective). The following parameters were used to quantify red objects and confluence: red (segmentation: adaptive, threshold: 1 RCU, edge split: on, edge sensitivity: -30, hole fill: 2 µm^2^, minimum area: 20 µm^2^, maximum area: 3000 µm^2^, maximum eccentricity: 1) and phase (segmentation: AI Confluence, hole fill: 10 µm^2^). To harvest, cells were placed on ice and each well was washed with 1 mL cold 1X PBS prior to the addition of 500 µL cold 1X MES buffer. Cells were lifted using a cell scraper and collected in 1.5 mL tubes. Samples were then sonicated using ten one second pulses and 60% amplitude with a Fisher Scientific Model 120 sonic dismembrator (Thermo Fisher Scientific). Samples were then centrifuged at 10,000 *x g* for 15 min at 4°C. Samples were then processed according to kit instructions. Briefly, samples were treated 1:1 with metaphosphoric acid (0.1 g/mL, Fisher Scientific, Cat# AC219221000), incubated at room temperature for 5 min, then centrifuged at 3,000 *x g* for 3 min at room temperature. Samples were transferred to a fresh tube and frozen at - 20C until needed. Immediately prior to performing the assay, samples were thawed on ice. 4M triethanolamine (abbreviated TEAM, Millipore Sigma, Cat# 90279-100ML) was then added to each sample at a ratio of 50 µL TEAM per 1 mL sample and vortexed briefly. Upon performance of the assay, absorbance was measured at 410 nm using a Synergy Neo2 plate reader (BioTek). Calculations were performed as described by the kit. Glutathione levels were normalized to the live cell density (mKate2^+^ objects/mm^2^) in each well immediately prior to cell harvesting.

#### STY-BODIPY analysis

On day 1, HT-1080 cells were seeded at a density of 1,000 cells per well of a 384-well plate, 5,000 cells per well of a 96-well plate, or 45,000 cells per well of a 12-well plate in phenol red-free growth medium (similar to the above described regular HT-1080 growth medium but with phenol-red free DMEM (Cat# 17-205-CV, Corning) supplemented with L-Glutamine (either directly or using a stock solution as described in the cystine deprivation experimental methods). On day 2, STY-BODIPY (Cat# 27089, Cayman Chemical Company) dissolved in L-Glutamine supplemented phenol red-free HT-1080 growth medium was added to the wells, such that the final concentration was 1 µM. Cells were incubated with STY-BODIPY for 20 min prior to removal of the medium. Cells were then treated with the appropriate lethal compounds and cell death inhibitors and imaged every 1 h for a total of 16 h with an additional scan after 0.5 h or every 4 h for a total of 24 h with an additional scan after 1 h. Imaging was performed using a Sartorius S3 live cell analysis instrument, as described above, using green and red acquisition times of 300 ms and 400 ms, respectively, as well as the 20X objective. Non-oxidized (red; Integrated Intensity Per mm² (RCU x µm²/mm²)) and oxidized (green; Integrated Intensity Per mm² (GCU x µm²/mm²)) signals from experiments conducted in 384- or 12-well plates were quantified using the S3 software as follows: green (segmentation: top-hat with radius 50 µm, threshold: 0.4 GCU, edge split: on, edge sensitivity: -10, hole fill: 100 µm^2^, maximum area: 2000-3200 µm^2^) and red (segmentation: surface fit, threshold: 0.08-0.36 RCU, edge split: on, edge sensitivity: -20, hold fill: 60 µm^2^, minimum area: 0-20 µm^2^, maximum area: unspecified or 4000 µm^2^). The ratio of the red:green signals was then calculated prior to plotting. For experiments conducted in a 384-well plate, wells with strongly fluorescent debris were omitted from the final analysis. For experiments conducted in a 96-well plate, the following parameters were used to detect red and green objects: green (segmentation: surface fit, threshold: 0.3 GCU, edge split: on, edge sensitivity: - 10, hole fill: 100 µm^2^, maximum area: 2000 µm^2^) and red (segmentation: surface fit, threshold: 0.15 RCU, edge split: on, edge sensitivity: -20, hole fill: 60 µm^2^, minimum area: 20 µm^2^, maximum area: 4000 µm^2^).

#### Amino acid analysis

HT-1080^N^ cells were seeded at a density of 2.4 × 10^5^ cells per well of a 6-well plate. The following day, wells were treated with DMSO (vehicle control), bortezomib (400 nM), or carfilzomib (400 nM) for 4 h. Each well was then washed twice with cold 1X PBS. Next, 500 µL cold 1X PBS was added to each well and cells were lifted from the plate using a cell scraper. The cell suspension was then transferred to a 1.5 mL Eppendorf tube, and cells were pelleted by centrifugation at 500 x *g* for 5 min at 4°C. PBS was aspirated and cell pellets were stored at -80°C. Amino acid analysis was performed at the Molecular Structure Facility Proteomics Core at the University of California at Davis as follows. Samples were resuspended in 200 µL 5/50 (5% formic acid/50% ACN). 100 µL of the cell solution was transferred to each of two glass hydrolysis tubes, then dried. To analyze Cys and Met, samples were oxidized in fresh performic acid at 2°C overnight then dried. All samples then underwent liquid phase hydrolysis (Pierce Sequanal grade 6N HCl, 1% phenol, 110°C, 24 h, in vacuo). Subsequently, samples were cooled, unsealed, and dried. Hydrolysates were then dissolved in a buffer consisting of Pickering and 40 nmol/mL NorLeucine (internal standard). 50 µL sample was then injected onto a Concise Ion-Exchange column. Amino acid analysis was performed using a Hitachi 8800 or 8900 analyzer, which were calibrated using amino acid standards compared to the NIST standard reference material 2389a in each sequence (Millipore Sigma, Cat# A9906).

#### Biochemical proteasome activity measurement

Chymotrypsin-, trypsin-, and caspase-like proteolytic activities were measured using the Proteasome-Glo 3-Substrate Cell-Based Assay System (Cat# G1180, Promega, Madison, WI). On day 1, HT-1080^N^ cells were harvested, pelleted by centrifugation at 350 x *g* for 4 min at room temperature, washed with HT-1080 complete growth medium, pelleted by centrifugation at 350 x *g* for 4 min at room temperature, and resuspended in HT-1080 complete growth medium. Cells were then seeded at a density of 4,000 cells per well of a 96-well white-walled assay plate (Corning, Cat# 3610). On day 2, cells were treated with the relevant compounds for 2 h. Epoxomicin (10 µM) was sometimes used as a positive control for proteasome inhibition. Immediately prior to performing the assay, a Sartorius S3 instrument was used to image the plate using the 10X objective with the red channel acquisition time set to 800 ms. Counts of mKate2-positive red objects (i.e., live cells) per mm^2^ were used for subsequent data normalization. Proteasome activity was then measured according to kit instructions. Briefly, substrate reagents were added to the assay plate and luminescence was measured using the following protocol: orbital shaking for 2 min at 800 rpm, 15 min delay, luminescence reading with gain of 100, integration time of 10 s, read height of 1 mm using a Synergy Neo2 plate reader (BioTek). Data were analyzed according to kit instructions and proteasome activity was normalized to live cell counts per well.

#### Imaging-based proteasome activity measurement

On day 1, HT-1080 or 293T cells stably expressing pLenti mCherry-T2A-eGFP were harvested, pelleted, and resuspended in L-Gln supplemented phenol red-free growth medium. HT-1080 or 293T cells were seeded at a density of 1,500 or 3,500 cells per well, respectively, in a 384-well black-walled assay plate (Corning, Cat# 3764). On day 2, cells were treated for 2 h with ten-point, two-fold dose responses of erastin2 (high dose 5 µM), RSL3 (high dose 5 µM), bortezomib (high dose 400 nM), carfilzomib (high dose 400 nM), or with the vehicle control DMSO in addition to ferrostatin-1 (1 µM) or the vehicle control DMSO. Following the 2 h treatment, cells were treated with cycloheximide (final concentration 178 µM) and imaged every 4 h for a total of 16 h using an S3 live cell analysis instrument using a 10X objective and with a red channel acquisition time of 800 ms and a green channel acquisition time of 400 ms. Red (mCherry) signal (Integrated Intensity/mm² (RCU x µm²/mm²)) and green (GFP) signal (Integrated Intensity/mm² (GCU x µm²/mm²)) were quantified using the S3 software with following metrics: green (segmentation: top-hat, radius: 50 µm, threshold: 0.95 GCU, edge split: on, edge sensitivity: -80, hold fill: 80 µm^2^, minimum area: 60 µm^2^) and red (segmentation: surface fit, threshold: 0.5 RCU, edge split: on, edge sensitivity: -50, hole fill: 60 µm^2^, minimum area: 60 µm^2^). The ratio of green:red signal was then calculated prior to data plotting.

### QUANTIFICATION AND STATISTICAL ANALYSIS

Google Sheets (Alphabet Inc., Mountain View, CA) and Microsoft Excel 16.101.1 (Microsoft Corporation, Redmond, WA) were used to conduct mathematical and statistical analyses, including computation of lethal fractions and Bliss independence model-based analyses. Data were plotted using Prism 10 (GraphPad Software, La Jolla, CA). Protein on western blots was quantified using EmpiriaStudio version 3.3.0.195 (LICORbio, Lincoln, NE). Adobe Illustrator 2025 (Adobe Systems, San Jose, CA) was used to create figures. R version 4.5.1 (R Foundation for Statistical Computing, Vienna, Austria) and RStudio version 2026.01.1+403 (Posit Software, Boston, MA) were used for some data analyses. Cytoscape version 3.10.4 (cytoscape.org) was used to generate the protein interaction map.

