## Supplemental Figure 1 for "Dissecting Complex Interactions Between Ferroptosis and the Proteasome"

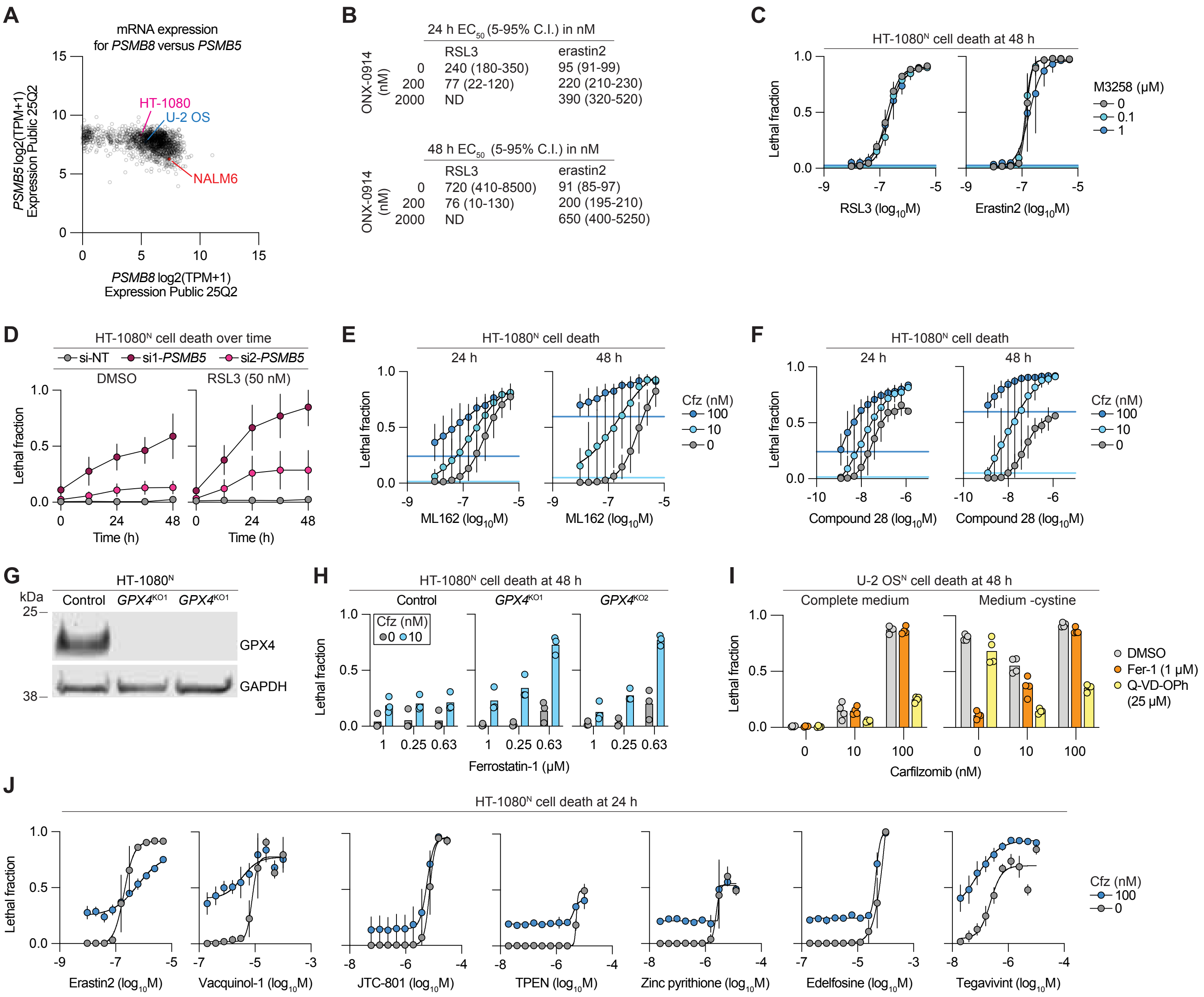


**Supplemental Figure 1: Ferroptosis regulation by the proteasome, related to Figure 1.** (A) Gene expression. From the DepMap database. Each datapoint represents one cell line. The position of three cell lines are indicated. (B) Quantification of cell death effects. Values are derived from analysis of the curves presented in main Figure 1. Results are from three independent experiments for RSL3 and two independent experiments for erastin2. (C) Cell death determined by imaging of live (nuclear mKate2-positive) and dead (SYTOX Green-positive) cells. Live and dead cell counts were integrated into the lethal fraction score (0 = all cells in the population alive, 1 = all cells in the population dead). (D) Cell death determined by imaging. SiRNA transfections were conducted 48 h prior to the start of compound treatments and cell death monitoring. Si-NT: non-targetting control siRNA. (E) Cell death determined by imaging. Cfz, carfilzomib. Results are from four independent experiments. (F) Cell death determined by imaging. Results are from three or four independent experiments per condition. (G) Protein expression determined by immunoblotting. Representative of three independent blots. (H) Cell death determined by imaging. Control and *GPX4*$^{KO1/2}$ cells were grown in medium containing ferrostatin-1 prior to the start of the experiment, then switched into medium containing different amounts of ferrostatin-1 ± carfilzomib. (I) Cell death determined by imaging. Results are from four independent experiments. (J) Cell death determined by imaging. Results in C,D and J represent mean ± SD from three independent experiments. Results in H and I are individual datapoints from independent experiments.
