## Supplemental Figure 2 for "Dissecting Complex Interactions Between Ferroptosis and the Proteasome"

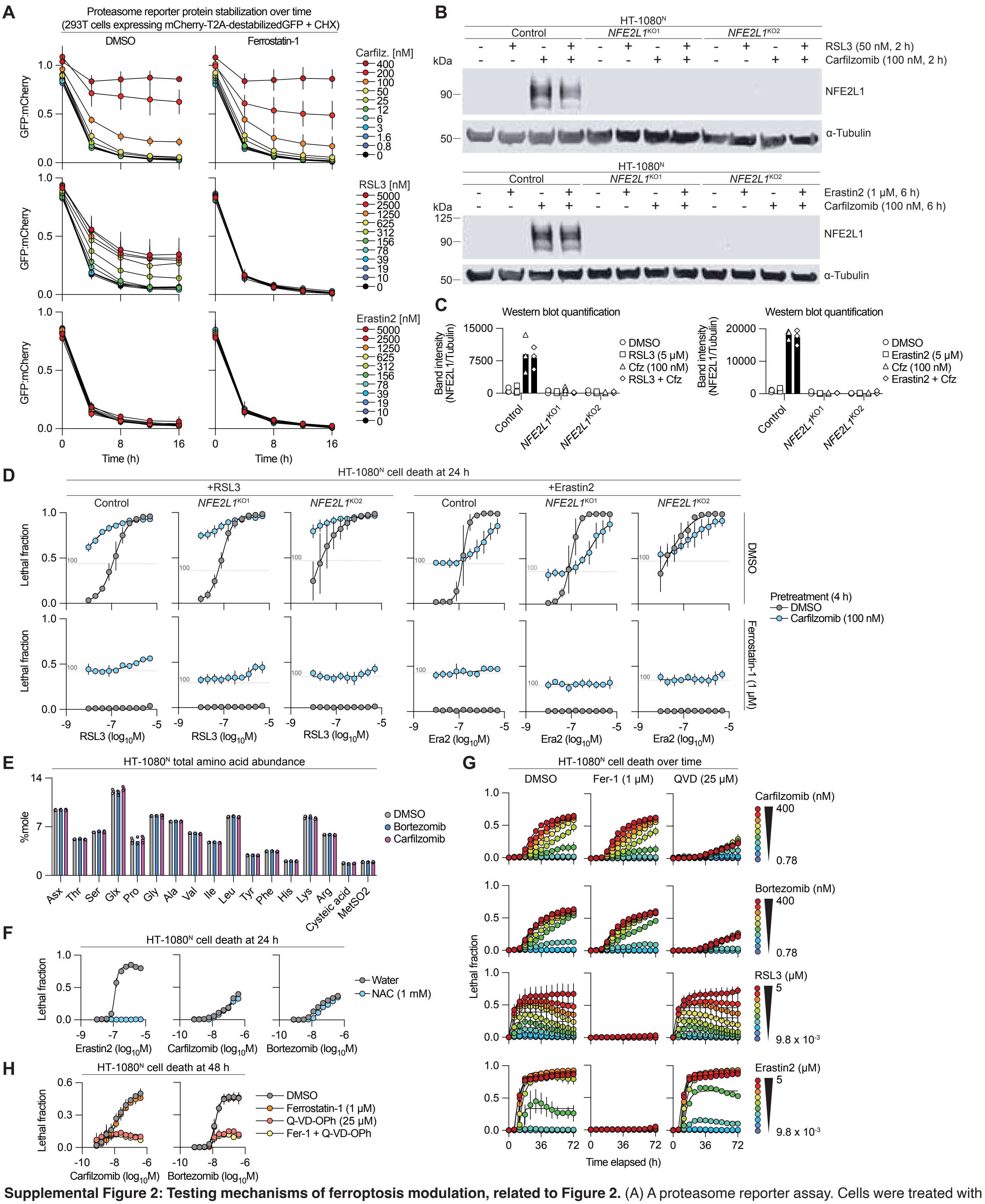


**Supplemental Figure 2: Testing mechanisms of ferroptosis modulation, related to Figure 2.** (A) A proteasome reporter assay. Cells were treated with inhibitors and cycloheximide for 2 h, then cycloheximide was washed out and cells were imaged for reporter protein stability. (B) Protein immunoblot. Representative of three independent blots. (C) quantification of results in (B). (D) Cell death determined by imaging of live (nuclear mKate2-positive) and dead (SYTOX Green-positive) cells. Live and dead cell counts were integrated into the lethal fraction score (0 = all cells in the population alive, 1 = all cells in the population dead). The label "100" indicates the basal cell death observed in the carfilzomib-alone condition, for reference. (E) Analysis of total cell amino acid content by mass spectrometry. Treatments were for 4 h. Carfilzomib and bortezomib were used at 400 nM. Asx = asparagine (Asn) + asparate (Asp). Cysteic acid = cysteine + cystine. (F) Cell death determined by imaging. (G) Cell death determined by imaging. (H) Cell death determined by imaging. Results in A, D, F, G and H are mean ± SD from three independent experiments. Results in C and E are individual datapoints from three experiments.
