## Supplemental Figure 3 for "Dissecting Complex Interactions Between Ferroptosis and the Proteasome"

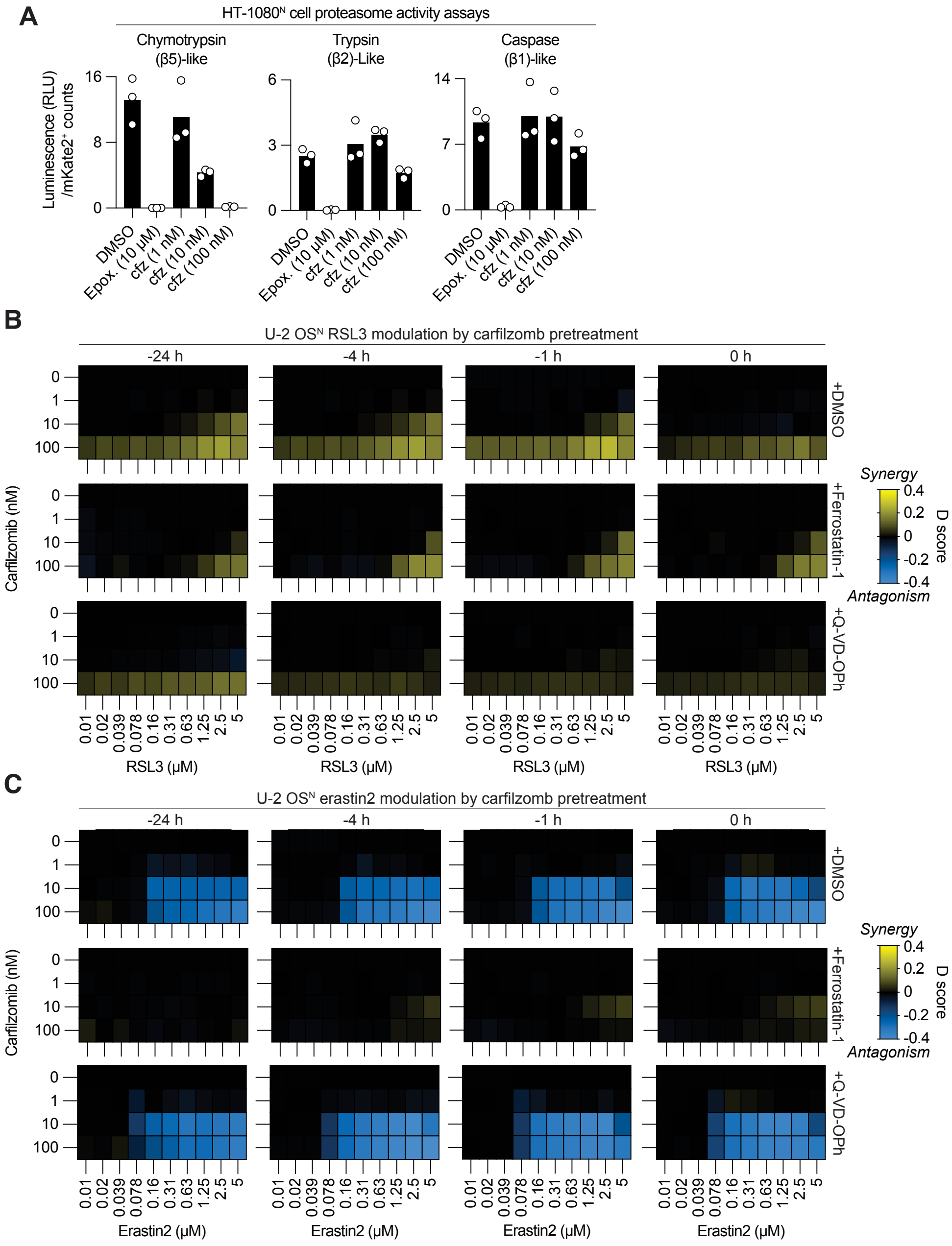


**Supplemental Figure 3: time-staggered kinetic modulatory profiling, related to Figure 3.** (A) Proteasome activity assays. Carfilzomib (cfz) treatmnt was for 90 min prior to the measurments. Epox., epoxomycin, a positive control proteasome inhibitor. Individual datapoints from seperate experiments are shown. (B,C) Summary of compound-compound deviation (D) scores. Ferrostatin-1 was used at 1 $\mu$M, Q-VD-OPh at 25 $\mu$M. Data are mean values from three independent experiments.
