## Supplemental Figure 4 for "Dissecting Complex Interactions Between Ferroptosis and the Proteasome"

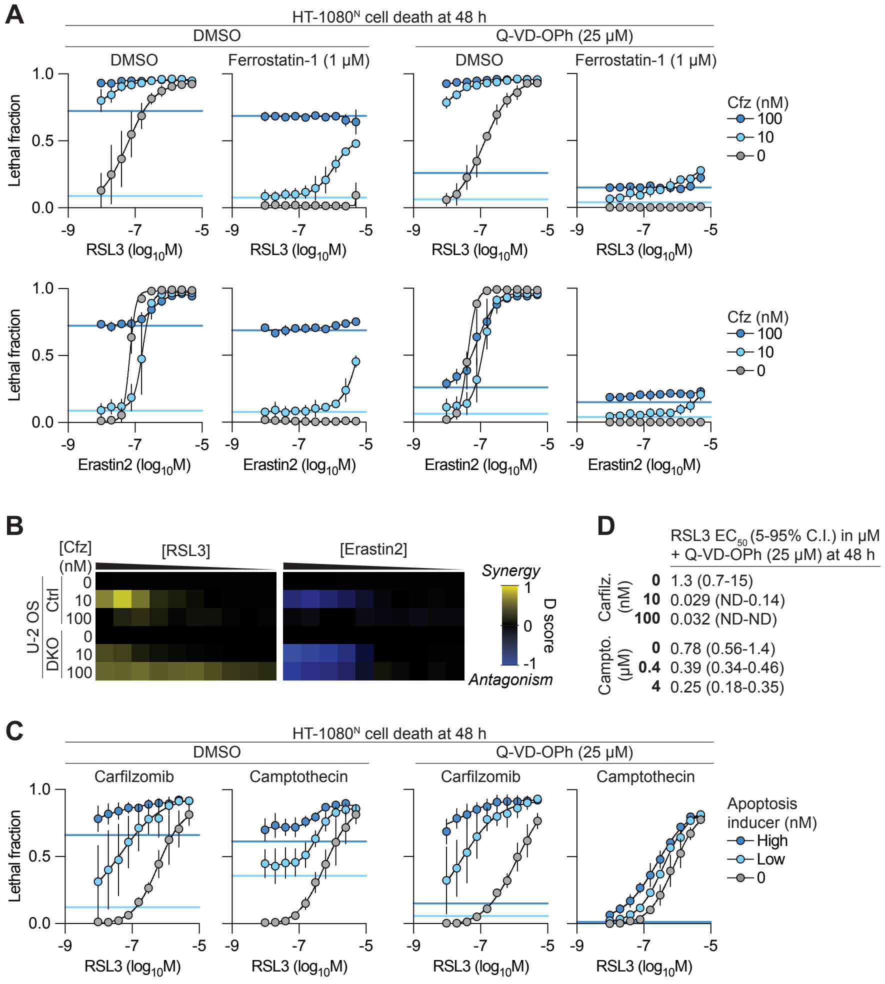


**Supplemental Figure 4: Ferroptosis execution independent of apoptosis, related to Figures 4.** (A) Cell death determined by imaging of live (nuclear mKate2-positive) and dead (SYTOX Green-positive) cells. Live and dead cell counts were integrated into the lethal fraction score (0 = all cells in the population alive, 1 = all cells in the population dead). (B) Summary of compound-compound deviation (D) scores. Heatmaps represent mean values from three independent experiments. (C) Cell death determined by imaging. High and low doses of carfilzimib were 100 and 10 nM, and high and low doses of camptothecin were 400 nM and 4 *μ*M. (D) Extracted parameter values for sigmoidal curve fits in (D). ND: not determined. Data in A, B and D represents mean ± SD from three independent experiments.
