## Supplemental Figure 5 for "Dissecting Complex Interactions Between Ferroptosis and the Proteasome"

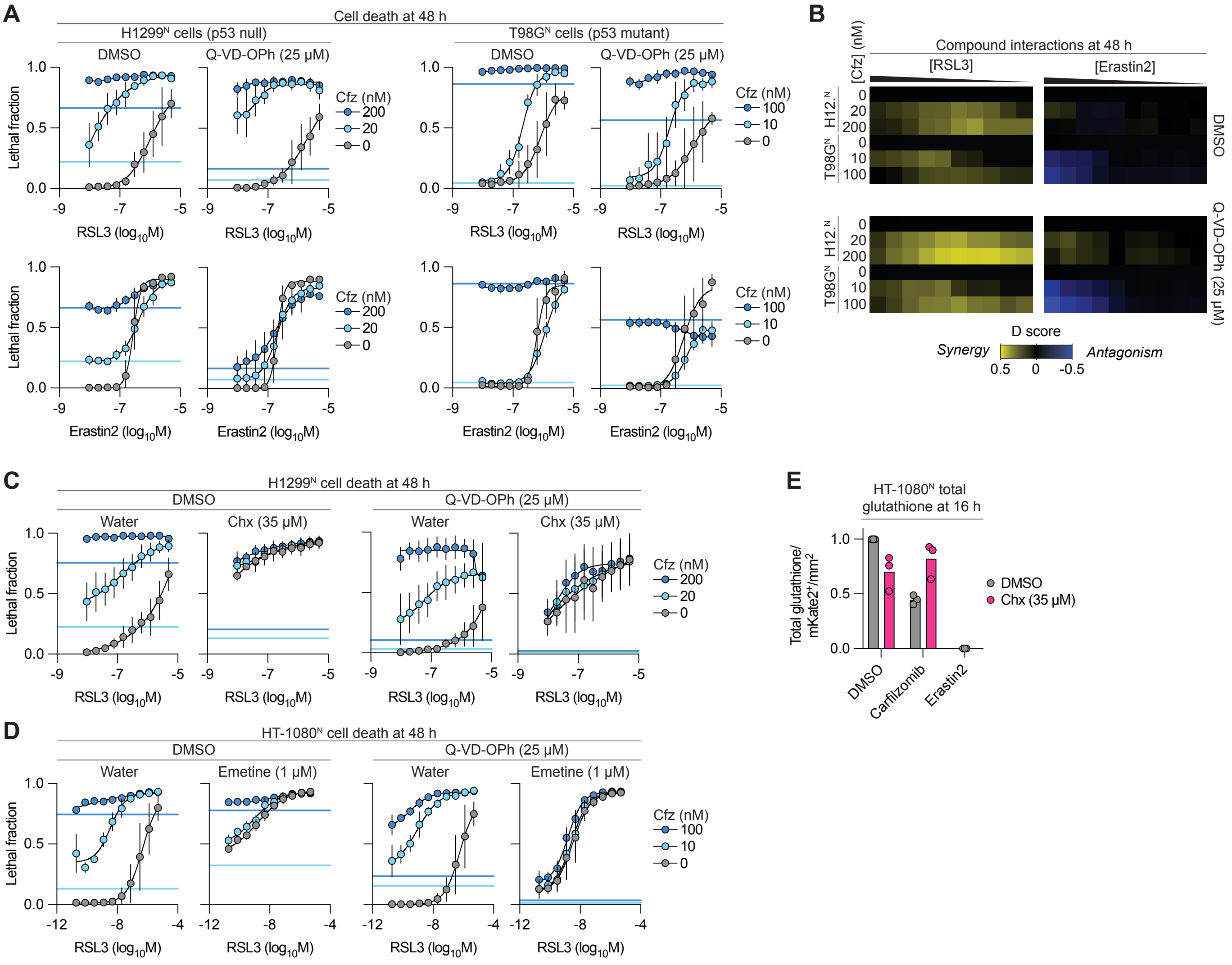


**Supplemental Figure 5: Protein synthesis promotes ferroptosis in response to proteasome inhibitors and GPX4 inhibition, related to Figure 5.** (A) Cell death determined by imaging of live (nuclear mKate2-positive) and dead (SYTOX Green-positive) cells. Live and dead cell counts were integrated into the lethal fraction score (0 = all cells in the population alive, 1 = all cells in the population dead). Cfz: carfilzomib. (B) Summary of compound-compound deviation (D) scores. Heatmaps represent mean values from three independent experiments. H12.: H1299. (D) Cell death determined by imaging. Chx: cycloheximide. (D) Cell death determined by imaging. All data represent mean ± SD from three independent experiments. (E) Total glutathione abundance determined using Ellman's reagent. Erastin2 is a positive control inducer of glutathione depletion. Individual datapoints from independent experiments are shown. Data in A, C and D represents mean ± SD from three independent experiments.
