## Supplemental Figure 6 for "Dissecting Complex Interactions Between Ferroptosis and the Proteasome"

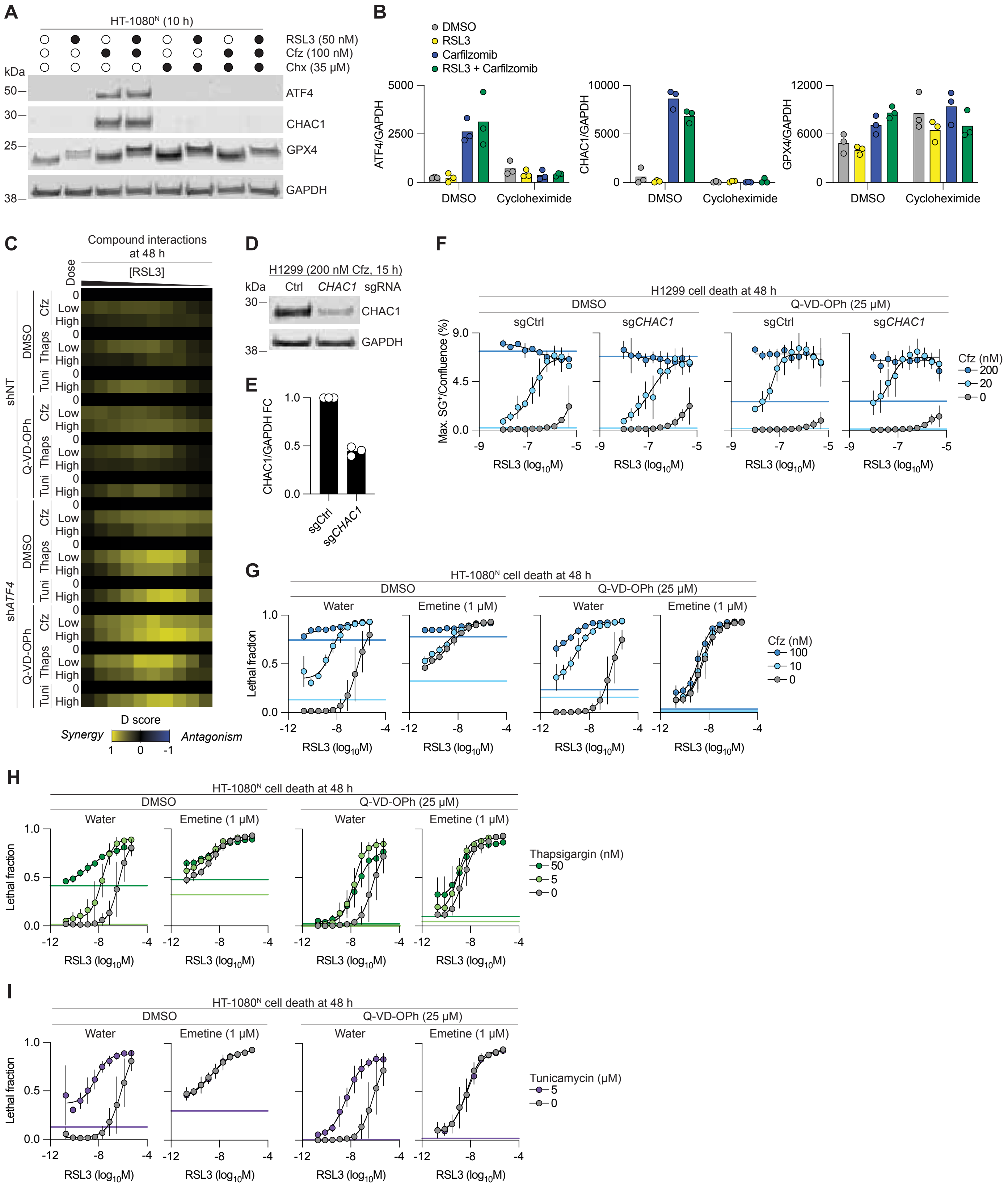


**Supplemental Figure 6: Modulation of ferroptosis sensitivity by ATF4 and other proteins, related to Figure 6.** (A) Protein immunoblot. Cfz: carfilzomib, Chx: cycloheximide. (B) quantification of results in A. (C) Compound interaction (deviation, D) analysis. Heatmaps represent mean values from three independent experiments. (D) Protein immunoblot. (E) quantification of results in C. (F) Cell death determined by imaging of dead (SYTOX Green-positive) cells. Dead cell counts were normalized to confluence at time = 0. (G) Cell death determined by imaging of live (nuclear mKate2-positive) and dead (SYTOX Green-positive) cells. Live and dead cell counts were integrated into the lethal fraction score (0 = all cells in the population alive, 1 = all cells in the population dead). Cfz: carfilzomib. (H) Cell death determined by imaging. (I) Cell death determined by imaging. Blots in A and C are representative of three independent experiments. Data in B and E are individual datapoints from seperate experiments. Data in F-I represent mean ± SD from three independent experiments.
